# A quiescent resident progenitor pool is the central organizer of tendon healing

**DOI:** 10.1101/2022.02.02.478533

**Authors:** Mor Grinstein, Stephanie L Tsai, Daniel Montoro, Heather L Dingwall, Ken Zou, Moshe Sade-Feldman, Miho J Tanaka, Terence D Capellini, Jayaraj Rajagopal, Jenna L Galloway

## Abstract

A tendon’s ordered extracellular matrix (ECM) is integral for transmitting force and highly prone to injury. How tendon cells, or tenocytes, embedded within this dense ECM mobilize and contribute to healing is unknown. Here, we identify a specialized *Axin2^+^* population in mouse and human tendons that remains latent in homeostasis yet initiates the healing response and serves as a major source of tendon progenitors. *Axin2^+^* tenocytes readily expand *in vitro* and express stem cell markers. *In vivo*, *Axin2^+^*-descendants are major functional contributors to repair: *Axin2^+^* tenocytes proliferate, acquire injury responsive states, and re-adopt a tenocyte fate post-injury. Specific loss of *Wnt* secretion in *Axin2^+^* cells alters their progenitor identity, disrupts their activation upon injury, and severely compromises any healing response. Our work highlights an unusual paradigm, wherein quiescent *Axin2*^+^ tenocytes self-regulate their identity and mobilization upon injury and provide the key initiating signal to organize a tendon-wide healing response.

---

Stem and progenitor cells rely on signaling interactions with their niche environment to maintain their state and regulate tissue maintenance and repair^1^. These signals can originate from neighboring cells, the ECM, daughter cells, or from the cells themselves^2–6^. Dense matrix tissues such as tendon present a unique constraint for progenitor cell regulation as there is a low cell density, limited turnover, and few cell-cell contacts. Yet despite these conditions, tendon cells can mobilize after injury to proliferate and repair the damaged tissue. Currently, the mechanisms underlying such regulation and the molecular identity of resident progenitor cell populations are poorly defined. Identifying central pathways and cell types capable of repairing tendon tissue *in vivo* would have major therapeutic implications as tendon injuries are a common clinical problem, accounting for up to 30-50% of all sports and work-related injuries^7^.

The tendon’s ECM consists of highly ordered type I collagen fibrils, which serve an important function in force transmission from the muscle to the bone. The hypocellular tendon is traditionally considered to contain two main populations of cells defined by their location and morphology: cells that surround the tendon in a sheath-like structure and elongated tenocytes with their long cellular projections within the tendon body^8^. Tenocytes reside within the ECM of the main tendon body and express the tendon transcription factor transgene, *Scleraxis* (*Scx*)-*GFP,* along with tendon matrix components including, *Collagen1a2* (*Col1a2*) and *Tenomodulin (Tnmd)* ^9^. Changes in matrix composition along the tendon length from the bone attachment, or enthesis, to the muscle attachment, or myotendinous junction, have been observed^10, 11^, and thought to result from differences in biomechanical and developmental factors specific to those regions^12–14^. Recent studies have revealed more tendon cell heterogeneity^15, 16^, than was previously appreciated but the function of distinct tendon cell subsets remains unclear.

Cells termed tendon-derived stem/progenitor cells (TDSPCs) were identified *ex vivo* through their characterization by surface marker expression, expansion, and ability to undergo serial transplantation into mice^17^, but their relationship to resident tendon cell populations *in vivo* is unknown. Previous analysis of radioisotope incorporation in human adult tendons have shown limited bulk tissue turnover after the age of adolescence^18^, making it unclear what role tendon stem or progenitor cells may have in maintaining adult tendon tissue. However, analysis of diseased tendon samples revealed increased tissue turnover rates^19^, and low levels of proliferation of *Scx*-*GFP^+^* cells has been detected throughout adulthood^20^. Furthermore, upon tendon injury, a robust cellular response occurs, which could be suggestive of stem or progenitor cell activity in the healing process. In identifying the origins of these contributing cell populations, previous lineage tracing studies have shown contribution of *Sma*^+^ and *Tppp3*^+^ cells from the surrounding paratenon sheath to tendon healing^15, 21, 22^, pointing to the presence of tendon stem or progenitor cells in the tendon sheath. However, other studies have shown major contributions to adult tendon healing from *Scx*-lineage cells^23, 24^. As *Scx* is considered a general marker of tenocytes that populate the main tendon body, it is unclear if stem cells reside within this dense collagen matrix and if so, what mechanisms, despite their ECM-rich environment, maintain their state and initiate their deployment in healing.

Here, we describe a latent yet injury-responsive *Axin2*^+^ tendon cell population that is regulated by their own Wnt secretion. Single cell RNA-sequencing analysis of adult tendons in homeostasis and healing reveals complex heterogeneity in tendon cell populations and underscores the central role of *Axin2*^+^ tenocytes in healing. In homeostasis, the *Axin2*^+^ cells reside in the main tendon body, have an elongated morphology with long processes, express *Scx*-*GFP*, and divide more slowly than their non-*Axin2* counterparts, but *ex vivo,* they readily expand and are enriched for the same markers found in TDSPCs. After tendon injury, the *Axin2*^+^ cells mount a robust injury response by down-regulating *Scx-GFP*, migrating to the wounded area, proliferating, and adopting a rounded cell morphology. During the healing process, they upregulate *Wnt* ligands and *Wnt* pathway components and eventually re-differentiate into tenocytes expressing characteristic tendon genes, including *Scx-GFP*, with elongated cellular morphology. We show that *Axin2*^+^ cells self-regulate their own *Wnt* ligand expression, progenitor identity, and activation upon injury via autocrine *Wnt* signaling. Furthermore, loss of *Wnt* signals specifically from *Axin2*^+^ cells severely compromise tendon healing, underscoring their key role in initiating the healing response. Thus, *Axin2*^+^ cells represent a quiescent tendon progenitor cell population that resides in the ECM-rich main tendon body and that self-regulate their maintenance and mobilization upon injury.

## Results

### *Axin2* marks a slowly dividing tendon progenitor cell population

The *Wnt* responsive lineage tracing mouse line, *Axin2:Cre^ERt2^; ROSA^LSLTdTomato^* (abbreviated hereafter as *Axin2^TdTom^*)^25, 26^, was found to label a subset of cells in postnatal and adult tendons. As *Axin2* marks stem and progenitor cells in other tissues, we sought to further define this cell population in the tendon. First, we examined if the number of *Axin2* cells changes from periods postnatal tendon growth to physiological homeostasis, and we found that the number of *Axin2^TdTom^* cells in the tendon decreases with age. Tamoxifen (TAM) treatment at neonatal stages on postnatal day (P) 2 when the tendon is growing longitudinally and has significant cell proliferation resulted in 28 ± 6% of the total cells being labelled by *Axin2^TdTom^* at P80 (Fig. 1A, C; Extended data Fig. 1-1 2G). Labelling at P60 during homeostasis when longitudinal tendon growth and cell proliferation has slowed^20^, showed that 10 ± 2% of the cells were *Axin2^TdTom^* at P80 (Fig. 1B, C; Extended data Fig. 1-2G). As no prospective antibodies can be used to specifically isolate tendon cells, our flow cytometry analysis was performed on total cells isolated from the tendon that had be negatively sorted for blood (CD45) and endothelial cells (CD31). Consistent with these *Axin2^TdTom^* observations, we found decreased expression of Wnt ligands and pathway genes, less accessible associated genomic regions over time, and reduced *Axin2* expression upon integrated analysis of RNA-seq and ATAC-seq datasets of tendon cells from P0 to P35 (Extended data Fig. 1-1A-B, Extended data Fig. 1-2 D, E). Together, this analysis shows that *Wnt* signaling decreases as the tendon shifts from periods of growth (P0-P21) to physiological homeostasis (>P35)^20^, and this correlates with the number of *Axin2^TdTom^* cells in the tendon. To verify that *Axin2^TdTom^* cells are primarily tenocytes from the main tendon body, we examined *Axin2^TdTom^* cells for expression of the tenocyte marker, *Scx*-*GFP*, by flow cytometry and 2-photon microscopy from 3-5 months of age. We found that a subset 10±2% of *Scx-GFP* cells were *Axin2^TdTom^*-positive and that double *Scx-GFP^+^*/ *Axin2^TdTom^* cells were found in the main tendon body (Fig. 1 E-H, Extended data Fig. 1-2 A, F, Extended data Fig. 1-3, Extended data Fig. 1-4). Therefore, *Axin2^TdTom^* marks a subset of *Scx-GFP* tendon cells and the number of *Axin2^TdTom^* tendon cells diminishes from birth to adulthood.

**Figure 1.**
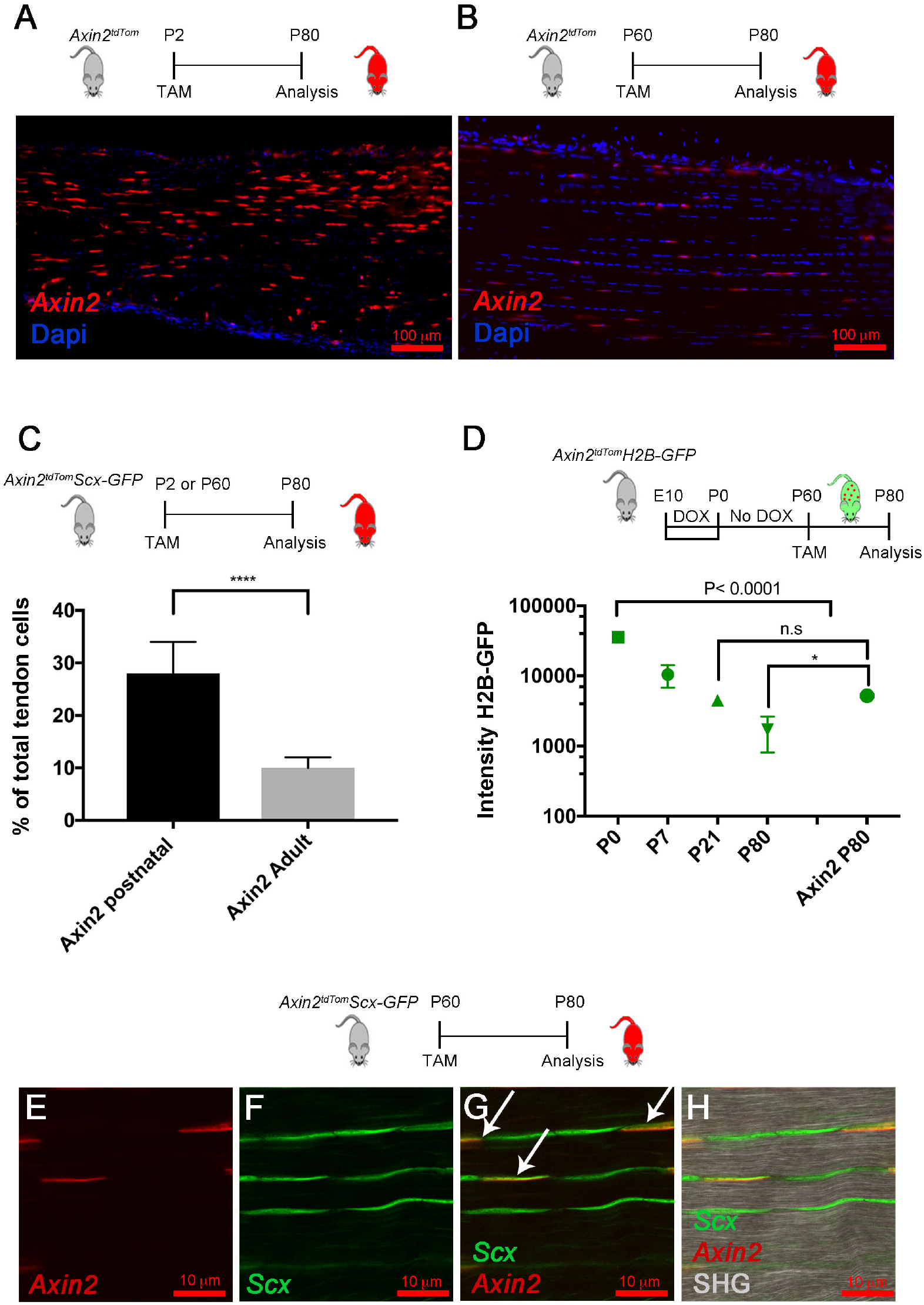
*Axin2^TdTom^* marks a latent subpopulation of tendon cells that decrease in number from birth. Tam was given to *Axin2^TdTom^* mice at postnatal day 2 (P2) or adults (P60, and the tissue was collected at P80. There are greater numbers of *Axin2^TdTom^* cells when labeled with Tam at P2 (A) compared with P60 (B) in Achilles tendon sections and quantified using flow cytometry at P80 (P2 cohort, n= 6 mice; Adult/P60 cohort, n= 8 mice, two-tailed unpaired T-test; ****p<0.0001). (C). The H2B-GFP system was used to evaluate tendon cell proliferation. Dox was administered to timed pregnant females from Embryonic day (E) 10 to birth, and at birth, Dox was removed. For *Axin2^TdTom^* labeling, Tam was given at P60 and cells were analyzed at P80. GFP intensity was measured in relation to GFP beads, which calibrates GFP intensity between experiments. We enriched for tendon cells by negatively sorting for CD34/CD45. *Axin2^TdTom^* cells had significantly higher H2B-GFP intensity compared to total tendon cells at P80 (n=4 mice per time point (P0, P7, P14, P21, P80, P80 *Axin2^TdTom^*, One-way ANOVA, F=148; n.s., not significant; *p<0.05) (D). (E-H) *Axin2^TdTom^* cells co-express *Scx-GFP* (arrows) in the main Achilles tendon body (20 Achilles tendons were examined by section) (SHG, second harmonic generation).

To define the proliferative activity of the *Axin2^TdTom^* cells during early postnatal and adult stages, we used the doxycycline (Dox) inducible Histone 2B-Green fluorescent protein (GFP) reporter mouse model (*Col1a1:tetO-H2B-GFP;ROSA-rtTA*), which can be used to quantify cell proliferation based on the dilution of H2B-GFP protein^27, 28^. After activating H2B-GFP expression by the addition of Dox from E10.5 to P0, we removed Dox at birth and allowed the cells to dilute their label based on their proliferation rates as previously reported^20^. This system is used to quantify cell proliferation and identify slowly cycling label retaining cell populations based on the stability and dilution of the H2B-GFP protein in each cell. We used GFP beads to calibrate measurements of GFP intensity decay between experiments. This allowed us to determine how postnatal *Axin2^TdTom^* tendon cells replicated since birth in comparison to non-*Axin2^TdTom^* cells. To enrich the sample for tenocytes, we examined H2B-GFP presence and intensity after excluding CD31^+^ endothelial cells and CD45^+^ blood cells. Surprisingly, we found that almost all the *Axin2^TdTom^* cells were H2B-GFP^+^ (95%), and we found that their intensity was significantly higher than that of non-*Axin2^TdTom^* cells with 5200±300 vs 1716± 904 fluorescence value at P80 (Fig. 1D). Taken together, this indicates that *Axin2^TdTom^* cells cycle less frequently than non-*Axin2^TdTom^* cells.

Tendon-derived stem/progenitor cells (TDSPCs) were characterized using cell culture and transplantation assays^17^. However, the identity and activity of the native cell population has been elusive. To examine whether adult *Axin2^TdTom^* cells display TDSPC characteristics *in vitro*, we Tam treated 3-month-old mice and harvested cells at 4 months. *Axin2^TdTom^* and non-*Axin2^TdTom^* cells were quantified by flow cytometry and next, cultured together for 10 days. Prior to cell culture, the *Axin2^TdTom^* cells comprised 9 ± 2% of the cell population and by day 10 of culture they were 47%± 5% of the cells (Fig. 2A-E), which represents an almost five-fold increase from their abundance in tendons. The *Axin2^TdTom^* cells changed morphology in culture, transitioning from thin spindle shaped cells with long cytoplasmic extensions to more rounded cells (Fig. 2A-C). To test if the *Axin2^TdTom^* cells express markers of previously described TDSPCs, we analyzed the surface marker expression of CD44, CD90.2, and Sca-1 by flow cytometry after 10 days in culture, but prior to passaging. We found that 97 ± 0.5% of *Axin2^TdTom^* cells expressed CD44, 89 ± 2.6% expressed CD90.2 and 71 ± 4% expressed Sca-1, indicating the *Axin2^TdTom^* cells are enriched for these surface markers (Fig. 2F-H), similar to the previously defined TDSPCs. The *Axin2^TdTom^* cells at day 10 in culture also were negative for CD31 and CD45 (Fig. 2I, J). To compare gene expression of *Axin2^TdTom^* and non-*Axin2^TdTom^* cells, the cells were harvested as described, but plated separately on day 0 of culture. After 10 days in culture, qRT-PCR analysis revealed that *Axin2^TdTom^* cells had increased levels of *Axin2* (Fig. 2K), suggestive of a persistent response to canonical *Wnt* signaling. We also observed greater relative expression of the tendon genes, *Scx*, *Mohawk* (*Mkx)*, and *Collagen 1a1*, and of *Ki67* (Fig. 2K), consistent with their tendon identity and ability to readily expand *in vitro*. Together, our results suggest the *Axin2^TdTom^* cells are the endogenous source of the previously described TDSPCs.

**Figure 2.**
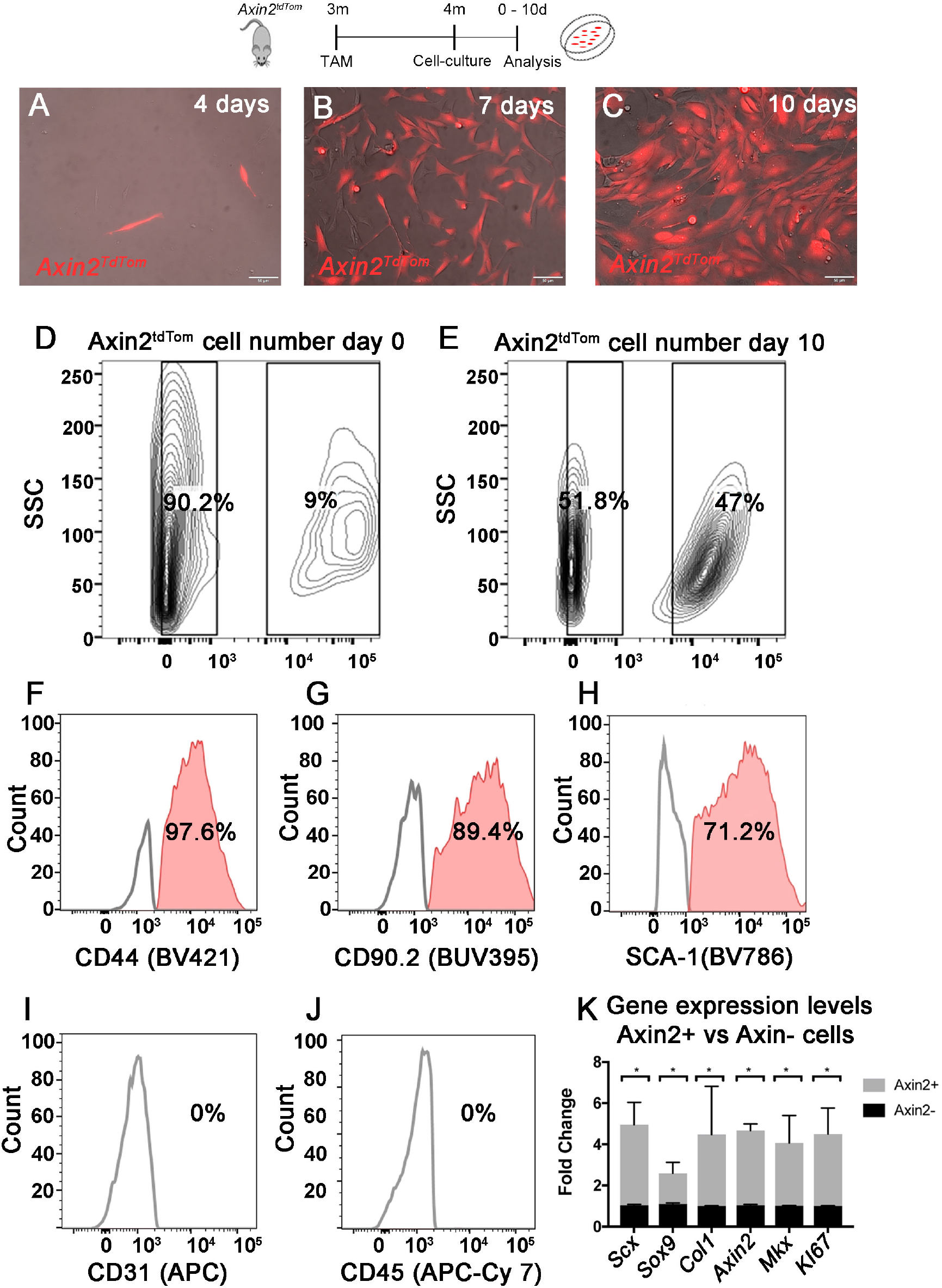
*Axin2^TdTom^* cells share characteristics with tendon-derived stem/progenitor cells. Tam was given to 3-month-old Axin*2^TdTom^* mice and at 4 months, cells were extracted from the limb tendons and analyzed or plated in cell culture. Images taken of *Axin2^TdTom^* cells show changing cell morphology at 4 (A), 7 (B), and 10 (C) days in culture. Representative flow cytometry plots showing *Axin2^TdTom^* and *Axin2^-^* cells at 0 and 10 days in culture (D-E). Flow cytometry histograms shows *Axin2^TdTom^* cells are enriched for CD44, CD90.2, and Sca-1 at day 10 in culture (F-H). Flow cytometry histogram shows *Axin2^TdTom^* cells do not express CD31 and CD45 after 10 days in culture (I-J). FACS was performed using limb tendon cells from n= 5 *Axin2^TdTom^* mice. *Axin2^TdTom^* cells express higher relative amounts of *Scx*, *Sox9*, *Col1a2*, *Axin2*, *Mkx* and *mKI67* transcripts by RT-qPCR compared with *Axin2^-^* cells after 10 days in culture (n=3 mice; Multiple T-test with two-stage step-up method of Benjamini, Krieger and Yekutieli; *p<0.05) (K).

### *Axin2^TdTom^* cells are major contributors to tendon healing

Although *Axin2^TdTom^* cells divide slowly *in vivo*, their ability to rapidly expand in culture led us to examine their behavior following injury. We treated 3-month-old mice with Tam to lineage trace *Axin2^TdTom^* cells and performed a partial acute excisional injury on the Achilles tendon at 4 months. Following injury, increased expression of several genes, including *Axin2*, *Scx,* and *Col1a2*, is observed by qRT-PCR in the healing Achilles tendon relative to uninjured contralateral controls (Extended data Fig. 3-1J). Using 2-Photon microscopy and second harmonic generation (SHG) imaging, we examined cell lineage and collagen organization and density in the healing tendon. At 10 days post injury (dpi), the wound site was infiltrated with *Axin2*-lineage cells and SHG signal was impaired (Fig. 3B, B’). *Scx*-lineage cells in *ScxCre^ERT2^; ROSA^LSLTdTomato^* mice (abbreviated as *Scx^TdTom^*) were also observed throughout the injury site at 10 dpi (Extended data Fig. 3-1B, B’), suggesting that the *Axin2^TdTom^* cells comprise a subset of the *Scx*-lineage population. This result is consistent with injuries in another tendon type^29^. At 10 dpi, *Axin2*- and *Scx*-lineage cells were not positive for *Scx-GFP* (Fig. 3B, B’, E; Extended data Fig. 3-1B, B’, Extended data Fig. 3-4), indicating *Scx-GFP* is downregulated upon injury. To determine if *Axin2^TdTom^* cells proliferate after injury, BrdU was administered during healing. We found that *Axin2^TdTom^* cells incorporate BrdU at the injury sites, but not in the contralateral uninjured tendons (Extended data Fig. 3-1 F, H, I). At 20 and 30 dpi, the *Axin2^TdTom^* cells co-expressed *Scx-GFP* and their cell morphology changed from rounded to elongated, indicative of differentiation into tenocytes (Fig. 3 C-D’, F, G, Extended data Fig. 3-5, Extended data Fig. 3-6). Similar to *Axin2^TdTom^* cells, *Scx^TdTom^* cells became elongated in appearance and expressed *Scx-GFP* after 20 dpi, (Extended data Fig. 3-1 C-D’). Quantification of cells in the injury site showed that *Axin2^TdTom^* cells comprise a major portion of the total cells in the healing region per defined area at 10, 20, and 30 dpi (Extended data Fig 4-1K; Fig 6G). Together, these data show that *Axin2^TdTom^* cells are the major cell population that infiltrates, proliferates, and differentiates into tenocytes after injury.

**Figure 3.**
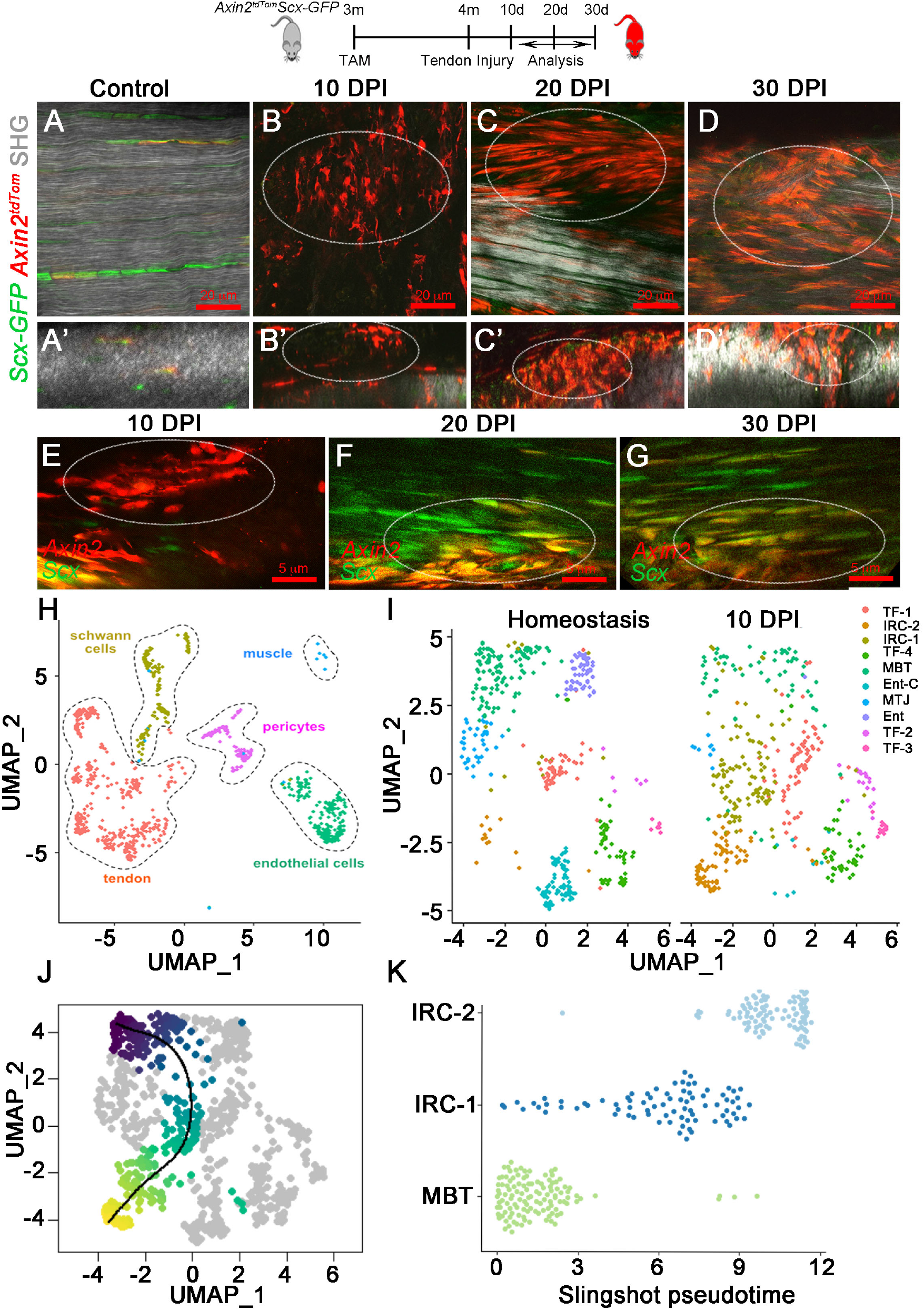
*Axin2^TdTom^* progenitor cells are major contributors to tendon healing. Tam was given at 3 months of age and Achilles tendon injury was performed at 4 months. 2-Photon microscopy images were taken of uninjured Achilles tendons from *Axin2^TdTom^* mice at 4 months (A, A’) and at 10, 20, and 30 days post-injury (DPI) (n>6 mice were examined at each time point). (B-D, circle indicates injured area) of the Achilles tendon of *Axin2^TdTom^; Scx-GFP* mice (A-D, sagittal views; A’-D’ transverse view resliced from sagittal sections of 0.4 μm). 2-Photon images show *Axin2^TdTom^* cells at 10 DPI at the injury site (E) and at 20 and 30 DPI co-expressing *Scx-GFP* (n>6 mice were examined for each time point). (F, G). UMAP representation of single cell RNA-seq data from Achilles tendons in homeostasis and 10 DPI (H). For homeostasis, 8 Achilles tendons were isolated from 4 adult mice at 4 months of age with Tam at 3 months. For 10 dpi, 3 Achilles tendons were harvested from 3 adult mice with tendon injury at 4 months of age. UMAP of sub-clustered tendon cells split by timepoint (I). Ten tendon clusters were identified: midbody tenocytes (MBT), myotendinous junction-associated tenocytes (MTJ), enthesis-associated tenocytes (Ent), entheseal cells (EnC), tendon cells 1-4 (TF1-4), and injury-responsive cells 1-2 (IRC1-2). Slingshot pseudotemporal reconstruction analysis inferred trajectory from the MBT to IRC-1 and IRC-2 clusters ((J, K).

### A single cell atlas of the adult tendon reveals activation of *Axin2^+^* mid-body tendon cells during healing

To further define *Axin2^TdTom^* cells in the broader context of all the cells in the tendon, we performed single cell RNA-sequencing during homeostasis and at 10 dpi (Fig. 3H-I). Mice were given Tam at 3 months; for homeostasis, cells were collected at 4 months and for 10 dpi, tendon injuries were performed at 4 months and cells collected after 10 days. Our analysis identified distinct tendon cell clusters. We also detected the presence of other cell types including Schwann cells, endothelial cells, skeletal muscle, and pericytes. Blood cells were present but removed from subsequent analysis. Clustering of tendon cells revealed distinct populations that we identified based on the enrichment of specific markers (Fig. 3H, extended Fig. 4-1A). The clusters include midbody tenocyte (MBT) (e.g. *Scx*+, *Tnmd*+, *Thbs4+*, *Fmod+*), myotendinous junction (MTJ)-associated tenocyte (e.g. *Col22a1*+, *Chodl, Scx*+)^30^, osteotendinous or enthesis-associated (Ent-C) tenocyte (e.g. *Pthlh*+, *Inhbb*+, *Scx*+), and entheseal (Ent; e.g. *Sox9*+, *Gli1*+, *Scx*-) cells in addition to four tendon fibroblast clusters (TF1-4) (Extended data Fig. 4-1A). TF1-4 did not express *Scx* (Fig. 5D) but these clusters were identified as tendon by their enriched expression of *Col1a1, Col3a1*, and other ECM components. Two additional clusters, identified as injury-responsive cell states (IRC-1, IRC-2), were found in homeostasis and expanded upon injury. IRC-1 and IRC-2 expressed genes known to be upregulated during tendon healing and associated with myofibroblast and mesenchymal identities, including *Acta2*, *Tagln*, *Sox9,* and *S100a4*^21^ (Fig 3I, Fig. 4A-F). Pseudotemporal reconstruction analysis of the UMAP representation using Slingshot^31^, inferred a trajectory from the MBT to IRC-1 and IRC-2 clusters (Fig. 3 J-K), suggesting that midbody tenocytes give rise to cells in these injury-responsive states.

**Figure 4.**
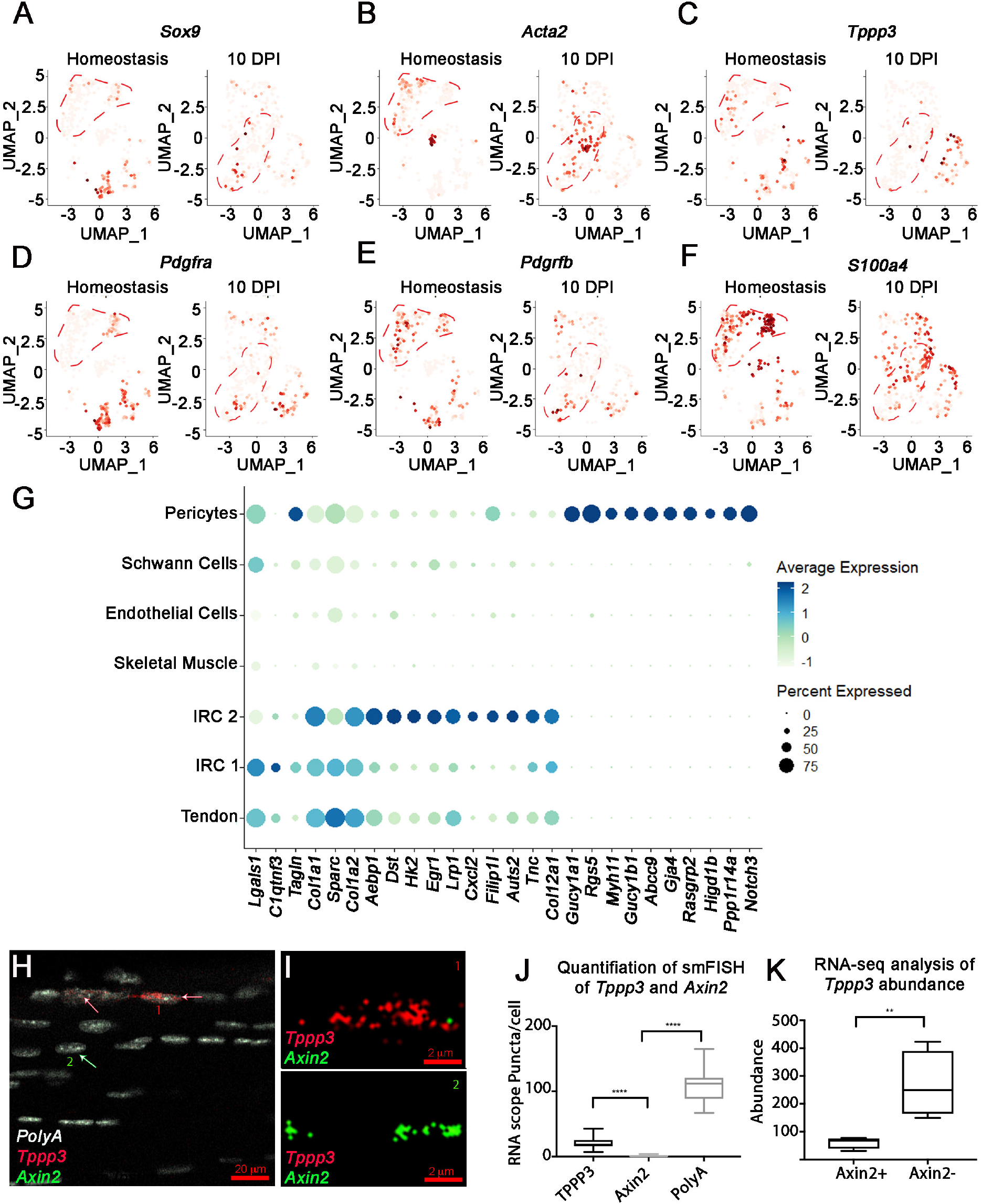
Gene expression in tendon clusters during homeostasis and injury. UMAP feature plots for different genes previously described to be involved in tendon healing during homeostasis and 10 DPI including *Sox9* (A)*, Acta2* (B)*, Tppp3* (C)*, Pdgfra* (D)*, Pdgfrb* (E), and *S100a4* (F). Tenocyte clusters (MBT, MTJ, and Ent) are outlined in red dotted lines for 0 dpi timepoint and injury-responsive cell states (IRC-1 and IRC-2) are outlined for 10 dpi timepoint (A-F). Dot plot showing gene expression in the pericyte, Schwann cell, endothelial cell, Skeletal muscle, IRC-1, IRC-2, and tendon clusters. This analysis shows similarities in tendon gene expression between IRC-1, IRC-2, and the tendon cluster (such as *TnC*, *Col12a1*, *Egr1*) and differences between IRC-1, IRC-2, and the pericyte cluster (such as *Notch3*, *Myh11*) (G) sMFISH of *Axin2* (green) and *Tppp3* (red) with *PolyA* (white) shows expression in different cells in sagittal sections of 4-month-old Achilles tendons (H). Magnification of two *Axin2* (green) and *Tppp3* (red) expressing cells (I). Quantification of *Axin2, Tppp3,* and *PolyA* puncta per cell shows most cells expressing *Tppp3* do not have *Axin2* puncta. *PolyA* was used as a staining control and to standardize measurements (n >20 cells from 2-3 sections from n=4 mice were quantified and statistical analysis was performed using two-way ANOVA (F=1.22); ****p<0.0001) (H-J). SMART-seq of *Axin2^TdTom^* vs non-*Axin2^-^*cells at 4 months of age show significant enrichment of *Tppp3* in the non-*Axin2* cells (n=4 mice; **p<0.01) (K).

**Figure 5.**
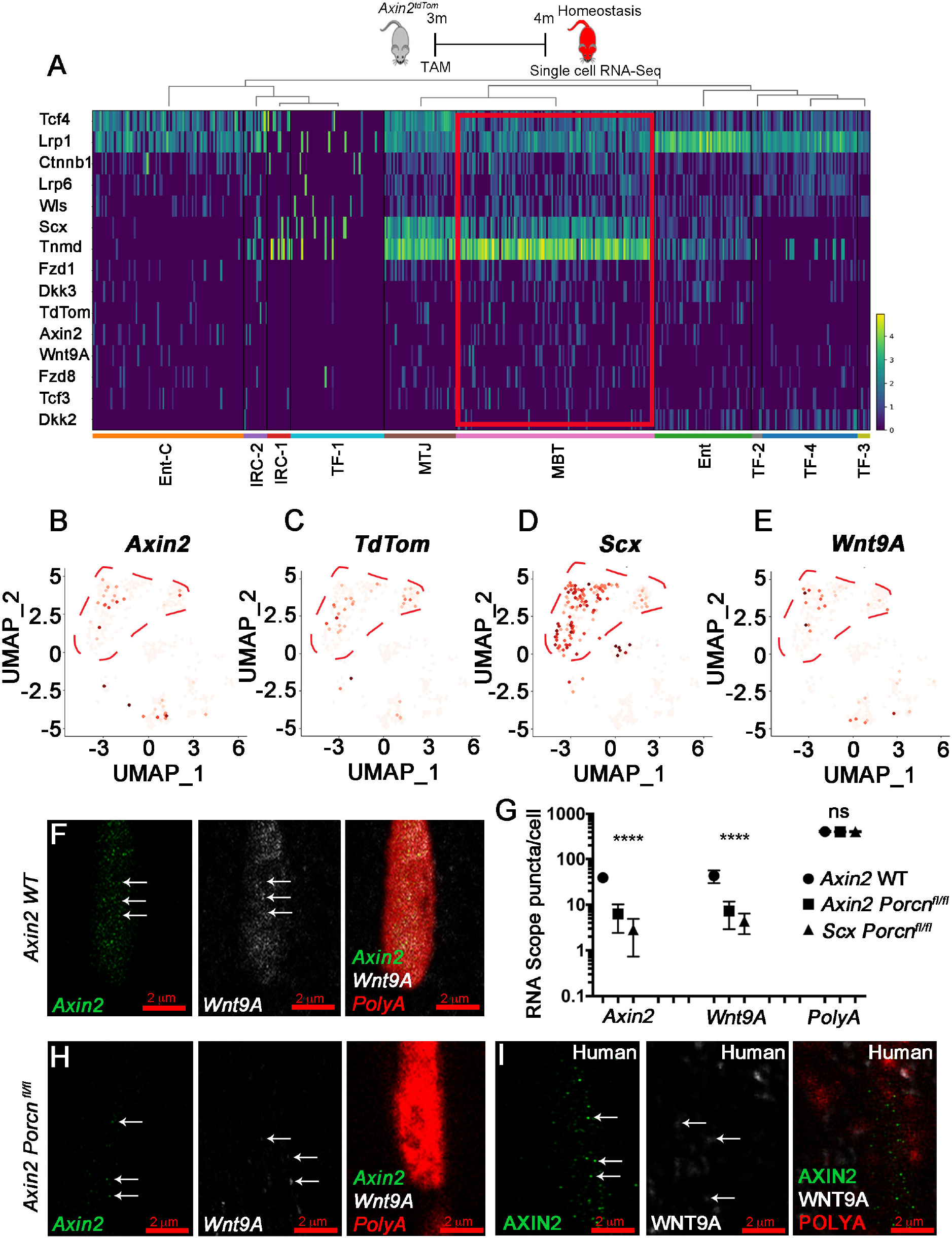
*Axin2^TdTom^* cells maintain identity in homeostasis through autocrine Wnt signaling. Single cell RNA-seq heatmap of canonical *Wnt* pathway components expressed in the different tendon clusters during homeostasis in 4-month-old *Axin2^TdTom^* mice; *Scx, Tnmd,* and *TdTom* expression is shown (A). UMAP feature plots of *Axin2, TdTom, Scx* and *Wnt9A* shows that these genes are expressed predominantly in the MBT cluster (B). smFISH images show one cell containing *Axin2* (green), *Wnt9a* (white), and *PolyA* (red) puncta in 4-month-old Achilles tendons from wild type (F) and *Axin2:Cre^ERT2^;Porcn^fl/fl^* mutants (H). Quantification of *Axin2*, *Wnt9a,* and *PolyA* puncta per cell in Achilles tendons shows co-expression of *Axin2* and *Wnt9A* in wild type and a decrease in puncta in *Axin2:Cre^ERT2^;Porcn^fl/fl^* mutants at 4 months of age (n >20 tendon cells were analyzed from each group with n> 8 Achilles tendon per group; groups consisted of 4 month old wild type, and *Axin2^TdTom^;Porcn^fl/fl^*, *Scx^TdTom^;Porcn^fl/fl^* with Tam given at 3 months; multiple T-test with Holm-Sidak correction was used for statistical analysis; n.s., not significant; ****p<0.0001) (G). *AXIN2* (green) and *WNT9A* (white) are co-expressed in human semitendinosus (hamstring) tendon cells (n>20 sections per sample, with n=4 separate samples,) (I). *PolyA* was used as an internal expression control (E-I).

To determine which cluster was enriched for *Axin2^+^* cells, we examined our single cell RNA-seq datasets for *Axin2* and *TdTomato* transcripts. *Axin2^+^* and *TdTomato*^+^ cells were predominantly found in the MBT cluster during homeostasis in concordance with our histological and 2-photon microscopy observations (Fig. 5A-C). *Axin2^+^* cells were also found in the enthesis clusters, Ent and Ent-C (Fig. 3I, 5B,C), which is consistent with a previous lineage tracing study of enthesis injury^32^. Although MTJ- and enthesis-associated tenocytes appear similar to the MBT, these distinct cell clusters should be located at the myotendinous and osteotendinous junctions, respectively, and not near the injury site, which is in the middle of the tendon body. Because of this and the presence of *Axin2* and *TdTomato*^+^ cells in the MBT, we focused our analysis on the MBT.

As our trajectory model indicates that MBT cells transition to IRC-1 and IRC-2 upon injury, we sought to test if *Axin2*-lineage cells express markers of IRC-1 and IRC-2 in healing. We focused on expression of *Sox9,* a gene expressed in skeletal progenitors and chondrocytes^33^, and *Acta2*, which encodes αSMA and is expressed in myofibroblasts as well as in growing and healing tendons^21, 34^. Both genes are expressed in IRC-1 and IRC-2 cells at 10 dpi (Fig. 4A, B). Immunofluorescent staining of injured Achilles tendons showed a majority of *Axin2^TdTom^* cells co-expressing Sox9 and αSMA at 10-30 dpi whereas uninjured tendons expressed limited amounts of Sox9 and αSMA (Extended data Fig. 4-1, Extended data Fig. 6-2B). As *Acta2* is highly associated with myofibroblasts or pericyte-like cells^35^, we examined the expression of additional genes to determine if IRC-1 and IRC-2 could represent a pericyte-like cell responding to injury rather than a tendon-derived cell. Analysis of gene expression in the different cell clusters after tendon injury shows that IRC-1 and IRC-2 share similarities in gene expression with the tendon clusters (*Egr1*, *TnC*, *Col12a1*) but do not express several markers that are enriched in pericytes, including *Notch3* and *Myh11* ( Fig 4 G).These results are consistent with *Axin2*-lineage tendon cells transitioning to IRC-1 and IRC-2 states in healing and supports a model in which *Axin2^+^* cells are specialized midbody tenocytes that are a major responding cell population in tendon injury.

To better define the *Axin2^TdTom^* cells before and after injury, we performed bulk RNA-seq analysis and compared *Axin2^TdTom^* to non-*Axin2^TdTom^* tendon cells during homeostasis and at 10 dpi. As the total RNA quantity in tendon cells is limiting and to avoid pooling samples or multiple tendon types which individually have a limited number of *Axin2^TdTom^* cells, we chose to amplify cohorts of pooled Achilles tendon cells using SMART-seq2 and perform differential expression analysis (Extended data Fig. 5-1 A, B) ^36^. During homeostasis, *Axin2^TdTom^* cells have significantly higher expression of several genes including *ProCR* (CD201) (Extended data Fig. 5-1 C), which marks stem/progenitor cell populations and is a target of Wnt signaling^37^. Immunostaining and flow cytometry analysis confirmed enriched expression of CD201 in *Axin2^TdTom^* cells (Extended data Fig. 5-1E). After injury, *Axin2^TdTom^* cells were significantly enriched for *Sox9,* consistent with our single cell RNA-sequencing and lineage tracing results, and for several tendon genes including *Col1a2, Tnmd*, and *Mkx* (Extended data Fig. 5-1 C). Of note, the previously described tendon sheath stem cell marker *Tppp3* was enriched in non-*Axin2^TdTom^* cells (lfc=-1.906, p value= 1.49e-9) (Extended data Fig. 5-1C), indicating *Axin2^TdTom^* cells are distinct from Tppp3^+^ cells, which have been shown to contribute to patellar tendon healing^15^. Examination of our single cell data set revealed *Tppp3* was expressed in several tendon cell clusters and did not appear restricted to a specific cluster (Fig. 4C). This result is consistent with a recent tendon single cell RNA-seq analysis^16^. Further expression analysis by small molecule FISH (smFISH) revealed *Tppp3*-expressing cells throughout the main tendon body and in the sheath (Fig 4H, I) but these cells did not significantly express *Axin2*, confirming our RNA-seq results. Therefore, *Axin2*^+^ cells represent a unique population of injury responsive cells that has yet to be defined (Fig 4 J,K).

### Autocrine Wnt regulation of *Axin2^TdTom^* cell identity

To interrogate how *Axin2^TdTom^* cells are regulated, we examined our RNA-seq datasets and as expected, we found enrichment of several canonical Wnt signaling components in the *Axin2*^+^ cell-containing MBT cluster (*Fzd1*, *Wnt9a, Dkk3*) (Fig. 5A) and enrichment for the Wnt pathway in *Axin2^TdTom^* cells (Extended data Fig. 5-1D). As *Axin2* is a direct target and negative regulator of canonical Wnt signaling^38^, these data are consistent with the notion that *Axin2*^+^ cells are regulated via canonical Wnt signaling. Intriguingly, we identified *Wnt9a,* a Wnt ligand that promotes the formation of synovial connective tissue cells ^39^, as enriched in the MBT cluster during homeostasis in our single cell data (Fig. 5A, E) and in *Axin2^TdTom^* cells in our bulk RNA-seq data (Extended data Fig. 5-1C). Using single molecule RNA fluorescent *in situ* hybridization, we confirmed that *Wnt9a* is expressed in *Axin2*^+^ cells (Fig. 5F). Therefore, we sought to test if *Wnt* signaling from the MBT *Axin2^TdTom^* cells themselves is required for their identity in homeostasis. As the MBT cluster is enriched for *Scx*+ cells, we first tested if *Wnt* signals originating from an intrinsic MBT population are necessary for *Axin2*^+^ cell identity. We deleted *Porcupine* (*Porcn*), a gene required for *Wnt* secretion, in *Scx*-expressing cells using *Scx:Cre^ERT2-TdTom^*; *Porcn^fl/fl^* mice by giving Tam at 3 months and examining gene expression by smFISH in sectioned tendons at 4 months. We found that *Axin2* and *Wnt9a* expression was decreased (Fig. 5G), suggesting that Wnt ligands originating from *Scx^+^* tendon cells are required to maintain *Axin2* expression during homeostasis. To specifically test if Wnt secretion from *Axin2^TdTom^* cells is necessary to maintain their identity, we Tam treated *Axin2-Cre^ERT2-TdTom^; Porcn^fl/fl^* mice at 3 months and examined their tendons at 4 months. We found decreased expression of both *Axin2* and *Wnt9a* upon *Porcn* deletion in *Axin2*-expressing cells (Fig. 5G, H). We confirmed that the cells were viable in *Porcn* mutants as we detected strong PolyA RNA signal, which also standardized our staining assay (Fig. 5 F-H). In addition, we found a decrease by flow cytometry in the percentage of *Axin2^TdTom^* tendon cells co-expressing *ProCR*/CD201 from *Axin2-Cre^ERT2-TdTom^; Porcn^fl/fl^* mice compared to wild type cells (Fig. S5-1 E-G). Despite *Axin2^TdTom^* cells comprising approximately 10% of the adult *Scx*^+^ cell population, no other cell source, including the *Scx*^+^ non-*Axin2^TdTom^* cells, could provide a Wnt signal to prevent the loss of *ProCR*, *Axin2*, and *Wnt9a* expression in *Axin2^TdTom^* cells lacking *Porcn*. Together, these data indicate that active Wnt secretion from *Axin2^TdTom^* tendon cells is required to maintain their cell state and *Wnt9a* expression, suggesting a positive feedback loop maintains their own identity.

Next, we wanted to determine if this unique *Axin2*+/*Wnt9a*+ cell is present in human tendons. We obtained healthy human hamstring tendons from patients ranging in age from 18 to 22 years and examined *AXIN2* and *WNT9A* expression. Within the main tendon body, we observed a subset of cells expressing *AXIN2* (11.7% ± 3.48 cells per total nuclei per 200 µm^2^ area), *WNT9A* (12.6% ± 4.7), and of which a majority of the *AXIN2+* cells co-expressed *WNT9A* (86.6% ± 7.6) (Fig. 5I). Together, these findings suggest that a similar *AXIN2+* cell population exists in human tendons.

### Loss of Wnt signaling from *Axin2^TdTom^* cells impairs healing

As loss of *Porcn* either in *Scx*- or *Axin2*-expressing cells negatively affects the ability of the *Axin2* cells to maintain their identity, we next tested if this loss affects their ability to respond to tendon injury. First, we deleted *Porcn* from the *Scx*-expressing cells (*Scx:Cre^ERT2TdTom^)* by giving Tam at 3 months and performing partial excisional tendon injuries at 4 months. Examination of *Scx:Cre^ERT2TdTom^*; *Porcn^fl/fl^* mice revealed a severely impaired healing response with virtually no *Scx*-lineage cells, reduced *Scx-GFP* expression, and a lack of matrix re-organization in the injured area at 30 dpi compared with wild type injured tendons (Fig. 6A, B, E; Extended data Fig. 6-1D; Extended data Fig. 6-2A). Unlike the control injured tendons, which display dynamic changes in the expression of tendon transcription factors (*Scx*, *Mkx*), matrix molecules (*Col1a2*, *Cola3a1)*, proteins found in mature tendons (*Fibromodulin, Connexin 43*^40^), proliferation (*MKi67)*, and genes associated with mesenchymal progenitors or differentiation towards other cell fates (*Sox9, Runx2*) (Extended data Fig. 3-1 J). *Scx:Cre^ERT2-TdTom^;Porcn^fl/fl^* injured Achilles tendons did not increase expression of many genes, including *Scx*, *Mkx Col1a2*, *MKi67*, and *Sox9*, at 10, 20, and 30 dpi (Extended data Fig. 6-1 A), suggesting a severely blunted healing response.

**Figure 6.**
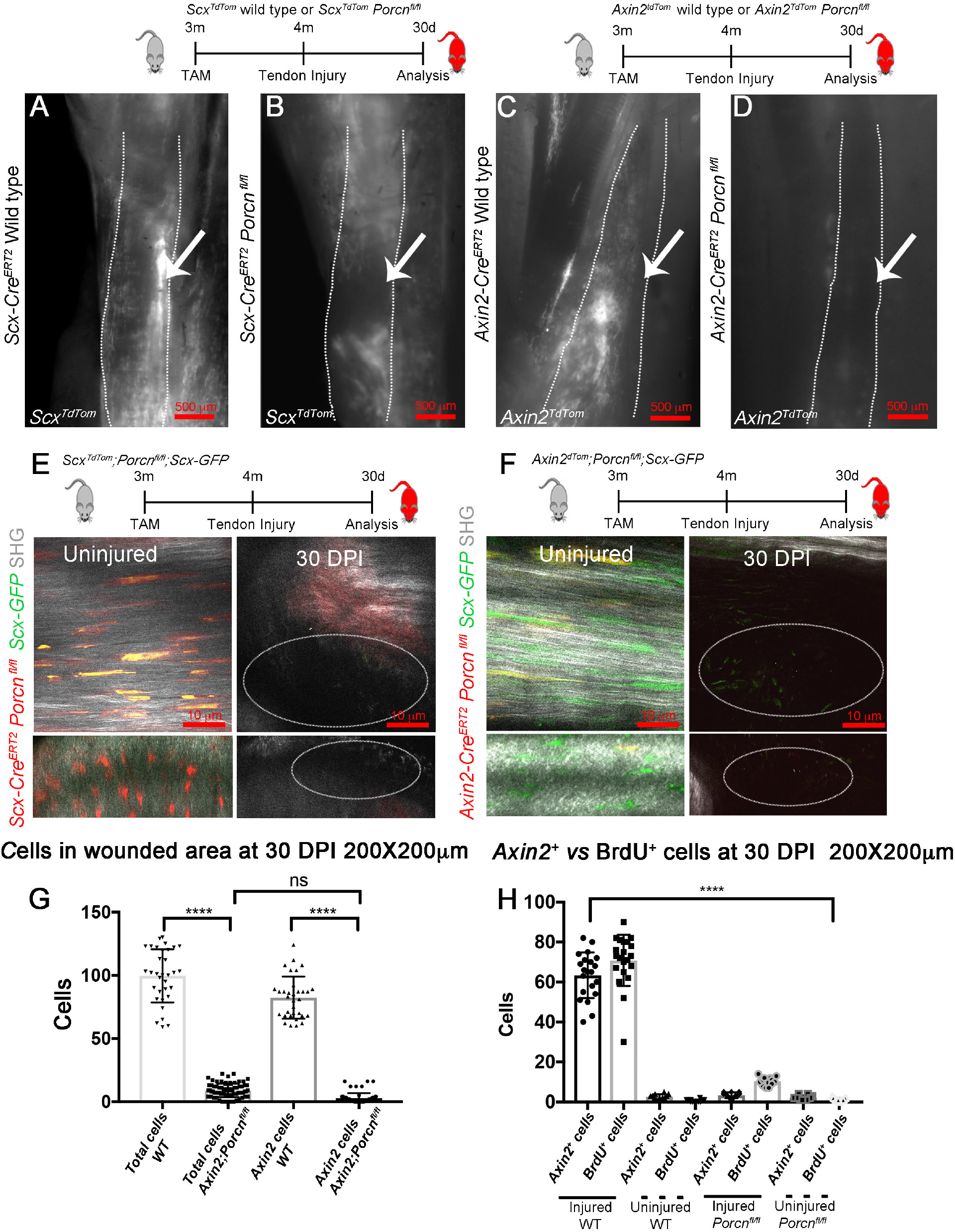
A cell-autonomous requirement for *Porcn* in *Axin2^TdTom^* cell injury response. Whole-mount fluorescent images of the Achilles tendon (outlined by dotted line in A-D) at 30 DPI of 5 months old mice that were given Tam at 3 months and injured at 4 months. *Scx^TdTom^* and *Axin2^TdTom^* cells localize to the injury site (arrows) in wild type controls (A, C), but not in *Scx:Cre^ERT2TdTom^;Porcn^fl/fl^* or *Axin2:Cre^ERT2TdTom^;Porcn^fl/fl^* (B, D). 2-Photon fluorescent and SHG images of uninjured Achilles tendons of 4-month-old mice (E, left) and injured 30 DPI *Scx:Cre^ERT2-TdTom^; Porcn^fl/fl^ ; Scx-GFP* Achilles tendons (E, right). 2-Photon fluorescent and SHG images of uninjured (F, left) and injured 30 DPI *Axin2-Cre^ERT2^; TdTom; Porcn^fl/fl^; Scx-GFP* (F, right) tendons of 5-month-old mice (Achilles tendons, n=6 mice per group per time point for A-F). Quantification of total cells and *Axin2^TdTom^* (Axin2 cells) cells in the injury site in wild type (WT) and in *Axin2-Cre^ERT2-TdTom^; Porcn^fl/fl^* mutants at 30 DPI (Achilles tendons from n=4 mice per group, 200X200 μM square area was quantified with 3 squares per section, n> 3 sections were analyzed per mouse per time point, Unpaired T test with Welch correction was used; n.s., not significant; ****p<0.0001) (G). Quantification of *Axin2^TdTom^* and BrdU+ cells in injured and uninjured Achilles tendons from *Axin2^TdTom^* wild type compared with *Axin2:Cre^ERT2-TdTom^;Porcn^fl/fl^* mutants at 30 DPI (Achilles tendons from n=5 mice per group with quantifications from 200 × 200μM square areas, with 3 areas per sections, and 3 sections per sample.. Unpaired T-test with Welch correction was used;; ****p<0.0001). (H).

**Figure 7.**
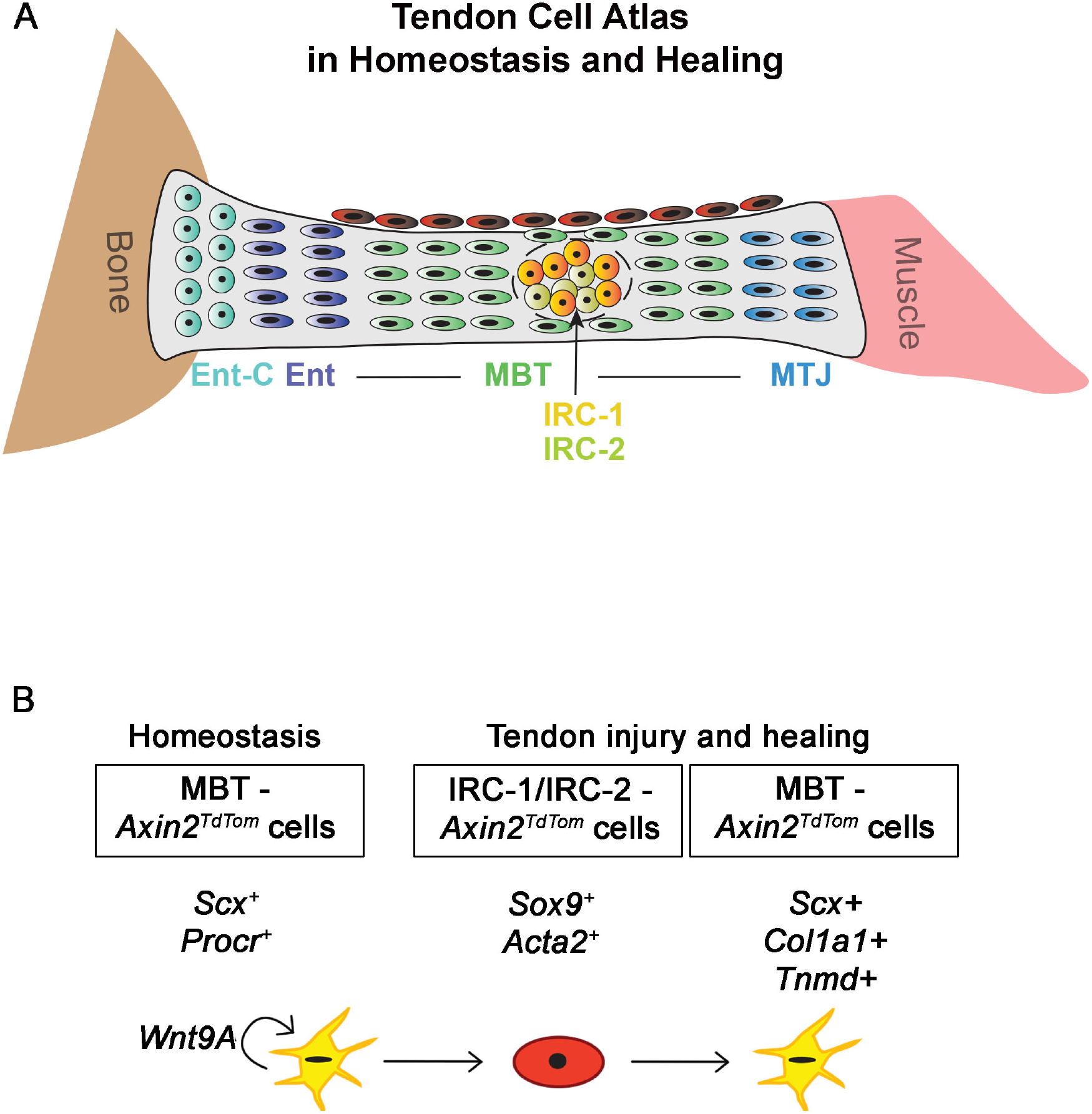
A single cell atlas of the mouse tendon illuminates the key role of *Axin2^+^* cells as central organizers of tendon healing. Cartoon depiction of the identified cell types in the tendon during in homeostasis and injury with a focus on relevant clusters (A). In our model (B) *Axin2^+^* progenitor cells reside within the mid-body tendon (MBT) cluster and require their own Wnt signal to maintain their Procr/CD201^+^ and *Axin2^+^* state in homeostasis. Upon injury, our single cell RNA-seq and genetic lineage tracing show that *Axin2*^+^ cells adopt an injury responsive state (IRC1, IRC2), expressing *Sox9* and *Acta2*. They robustly migrate, proliferate, adopt a rounded morphology, and then re-differentiate into tenocytes expressing common markers of tendon fate (*Scx*, *Col1a1*, *Tnmd*). These processes are entirely dependent upon secretion of their own Wnt ligands, likely *Wnt9a*.

To test if Wnt secretion specifically from the *Axin2+* cells is required for tendon healing, we analyzed *Axin2:Cre^ERT2-TdTom^;Porcn^fl/fl^* Achilles tendons at several stages after injury using the same Tam treatment and injury regimen. Similar to *Scx:Cre^ERT2-TdTom^;Porcn^fl/fl^* injured tendons, the healing response was significantly affected (Fig 6C, D, F; Extended data Fig. 6-1C) with disruption to the SHG signal, reduced cell proliferation, and abnormal morphology in healing *Axin2:Cre^ERT2-TdTom^;Porcn^fl/fl^* Achilles tendons compared to controls at 30 dpi (Fig. 6F,G,H; Fig. 3D. Extended data Fig. 6-2A,). Within the injury site, we observed a virtual loss of all *Axin2^TdTom^* cells as well as significantly fewer total cells and cells incorporating BrdU in *Axin2:Cre^ERT2-TdTom^;Porcn^fl/fl^* tendons compared to injured controls (Fig. 6C, D, F, G, H). Further examination revealed an increase in CD45^+^ cells at the healing site of *Axin2:Cre^ERT2-TdTom^;Porcn^fl/fl^* tendons compared to wild type tendons at 30 dpi (; Extended data Fig 6-2D,E,), suggesting that some of the cells present in *Porcn* mutants are blood cells. In addition, there was no significant increase in *Scx*, *Mkx Col1a2*, *MKi67*, *Fmod*, and *Sox9* expression along with several other genes in the injured compared to contralateral uninjured Achilles tendons of *Axin2:Cre^ERT2 TdTom^;Porcn^fl/fl^* mice (Extended data Fig. 6-1 B). Similarly, we observed significantly reduced numbers of *Axin2^TdTom^* cells that expressed Sox9 and αSMA in healing *Axin2:Cre^ERT2-TdTom^;Porcn^fl/fl^* Achilles tendons compared to controls (Extended data Fig 6-2B, C), suggesting the IRC-1 and IRC-2 clusters are not present in *Porcn* mutants. The similarities in the severity of the healing phenotype observed in the *Scx-* and *Axin2-*mediated *Porcn* conditional knock-outs supports a model in which *Axin2^TdTom^* cells are the major Wnt secreting subset of the *Scx* cells. In addition, the phenotype resulting from the specific deletion of *Porcn* in the *Axin2^TdTom^* cells strongly suggests that *Axin2^TdTom^* cells regulate their injury response in an autocrine manner. As the *Axin2^TdTom^* cell identity is affected in *Porcn* mutants, we cannot rule out that their inability to mount a healing response results from a loss of competence to respond to injury. Moreover, the lack of other non-*Axin2^TdTom^* cell populations to respond in injured *Axin2:Cre^ERT2^;Porcn^fl/fl^* indicates the *Axin2^TdTom^* cells through their secretion of Wnt signals are central to the recruitment of virtually all tendon cells in healing.

## Discussion

Our work identifies a unique *Axin2^+^* progenitor cell population that resides in the adult tendon, mobilizes upon injury to promote healing, and serves as the central organizer of the healing response. Our single cell analysis reveals unprecedented heterogeneity in tendon cell populations and positions the *Axin2^+^* mid-body tendon cells as key responders to tendon injury through regulation by their own Wnt signals. *Axin2^+^* cells are enriched for previously described markers of progenitor cell populations including ProcR/CD201, and expand extensively *in vitro*, while at the same time maintaining higher expression of markers of tendon fate. During homeostasis *in vivo*, *Axin2^+^* cells are latent, dividing less frequently than their non-*Axin2^TdTom^* counterparts, yet are poised to respond to injury, a characteristic of several stem and progenitor cell populations^41^. Consistent with this concept, *Axin2^TdTom^* cells are the major cell population responding to tendon injury. They infiltrate the injury site, proliferate, and re-differentiate into elongated *Scx-GFP* tenocytes. Therefore, we propose that *Axin2^TdTom^* cells are the endogenous cell source of the previously described TDSPCs ^17^.

Transcriptomics, single molecule *in situ* hybridization, and functional deletion of *Porcn* specifically in *Axin2^TdTom^* cells revealed that *Axin2^TdTom^* cells rely on their own secreted Wnt ligands to maintain their identity and injury response. The importance of Wnt secretion from the *Axin2^TdTom^* cells to maintain *Axin2*^+^ cell state in homeostasis is underscored by their loss not only of ProcR expression, but also of both *Axin2* and *Wnt9a* upon *Axin2:Cre^ERT2^*-mediated deletion of *Porcn*. This result suggests that the secretion of Wnt by *Axin2* cells is simultaneously maintaining *Axin2* identity and the expression of its own Wnt ligand, thereby providing a positive feedback loop to ensure the maintenance of the cell type. Autocrine regulation is a mechanism employed by stem and progenitor cell populations^42, 43^. As Wnt ligands act as short-range signals^44^, the *Axin2^+^* cell provides its own niche-like factors that act as a self-sustaining feedback loop. Autocrine signaling is uniquely suited for tendon progenitor cells residing in the main tendon body since they are embedded in a dense ECM with limited cell contacts.

Our work also positions *Axin2^+^* cells as key early initiators of tendon healing through their secretion of Wnt. We detect *Wnt9*a expression in *Axin2^+^* MBT cells during homeostasis (Fig. 5), and upon injury, our bulk RNA-seq shows significant upregulation of *Wnt9*a compared with non-*Axin2* cells (Extended data Fig 5-1), indicating *Axin2^+^* cells increase *Wnt9a* expression during a normal injury response. Consistent with this result, we observe increased *Axin2* transcript levels following injury (Extended data Fig. 3-1J). It is interesting to note that reports in the liver reveal alternative modes of regulation after injury in which there are no specialized subset of progenitors or stem cells. Instead, all hepatocytes upregulate *Axin2* and proliferate upon injury^45–47^. In contrast to this scenario, our results favor a model in which the *Axin2^TdTomato^* cells are a pre-defined subset of the *Scx^+^* MBT population. *Axin2^TdTomato^* cells are distinct from non-*Axin2* cells in their proliferation activity since birth (Fig. 1D), suggesting they are uniquely regulated. In addition, specific deletion of *Porcn* in the *Axin2^TdTomato^* cells 1 month prior to the injury dramatically affects their recruitment and proliferation during healing. There also are fewer non-*Axin2^+^* cells and an increase in CD45^+^ cells at the injury site, indicating loss of *Porcn* in the *Axin2^TdTomato^* cells more broadly affects the healing response. As the *Axin2^+^* cells represent a minority of the total tendon cell population, approximately 10% of the total tendon cells (Fig. 1C), the fact that the remaining non-*Axin2* tendon cells cannot compensate by secreting additional Wnt signals underscores the importance of the *Axin2^+^* cells in initiating the healing response. These data also suggest that non-*Axin2* cells cannot acquire *Axin2^+^* identity from other Wnt sources upon injury. How the adult *Axin2^+^* cells acquire their identity is currently unclear. Our transcriptomic and genomic analysis of the first postnatal month combined with our lineage tracing at P2 (Fig. 1A-C; Extended data Fig. 1-1), show increased Wnt pathway components and increased *Axin2^+^* cells, respectively, at early neonatal stages that decrease with age. This decline in Wnt signaling and *Axin2^+^* cells over time occurs concomitantly with a decrease in tendon cell proliferation^48^; marked matrix expansion^49^, and a loss of regenerative healing^34, 50^. It is possible that the adult *Axin2^+^* cells are descendants of a subset of these earlier *Axin2^+^* cells that remain latent yet retain the potential for regenerative healing in the adult. Whether *Axin2^+^* cell number continues to decrease at later stages is currently unknown, but it would be interesting to investigate if changes in *Axin2^+^* cell number or activity underlies age-related increases in tendon injury and disease.

Previous studies have shown that stem and progenitor cells from the tendon sheath are involved in tendon healing^15, 21, 22^. In contrast, we provide evidence for another cell population, marked by *Axin2*, that act as quiescent progenitor cells capable of activating upon injury to directly orchestrate the healing response. As *Scx* is expressed in tenocytes in the main tendon body, several lines of evidence support *Axin2* marking a subset of these *Scx^+^* MBT cells, including our bulk and single cell RNA-seq data and the similarities in lineage tracing and in phenotypes resulting from *Porcn* deletion using *Scx^CreERT2TdTom^* vs *Axin2^CreERT2TdTom^*. Further, previous studies have shown contribution from *Scx*-lineage and *Scx-GFP^+^* cells to tendon injury^23, 29^. We do observe non-*Axin2^TdTom^* cells at the injury site, and we speculate that these cells originate from the sheath. However, the lack of sheath specific markers complicates their definitive identification. Similar to another Achilles tendon single cell sequencing study^16^, we found that expression of *Tppp3*, a previously described sheath marker, was not restricted to a specific tendon cluster and was expressed in both the sheath and the main body of the Achilles tendon (Fig 4D, H-J), making it difficult to resolve which cluster may represent the sheath stem cell population. As the prior work also described these cells as *Tppp3*^+^ and *Pdfgra^+^*^15^, we examined the clusters and found co-expression of *Tppp3* and *Pdgfra* in a subset of clusters, Ent-C, TF-2, TF-3, and TF-4 (Fig 4D, E). It is possible that one of these clusters, particularly TF-4, which significantly expresses both genes, represents the previously described sheath population. Nevertheless, our expression analyses show that *Axin2^TdTom^* cells are distinct from *Tppp3*-expressing cells and our *Porcn* functional data positions *Axin2^TdTom^* cells as providing an upstream signal necessary for cell recruitment and proliferation at the injury site.

The identification of similar *AXIN2^+^/WNT9A^+^* cells in human tendons opens new potential therapeutic avenues to treat tendon disease and injury. Notably, the percentage of *AXIN2^+^* cells in humans is similar to adult mouse tendons, representing approximately 10% of the cell population. As we used human hamstring tendons for this analysis, it suggests *AXIN2^+^* cells may be broadly present in other tendon types. It is particularly interesting that *Axin2^+^* cells are enriched for *Wnt9a* in humans and mice as prior work has shown that this and other Wnt ligands have important roles in both promoting synovial connective tissue formation and suppressing chondrogenic potential in the context of in joint formation^39^. Wnt signaling also functions in cartilage and bone formation^51, 52^. As the appearance of ectopic bone and cartilage is associated with adult tendon overuse and injury^34^, it is interesting to speculate that perhaps aberrant regulation of Wnt signaling by the *Axin2^+^* cells could underlie these pathologies. Not only do *Axin2^+^* cells represent a new potential source for tissue engineering strategies, but also an exciting novel therapeutic target for tendon injury and disease. Therefore, we believe that our work identifying *Axin2^+^* cells as central organizers of tendon repair and their unique mode of self-regulation, together, will significantly impact our understanding of tendon healing mechanisms and the development of novel treatments for tendon injuries.

## Acknowledgements

We would like to thank Dr. Radhika Khetani and the Harvard T.H Chan School of Public Health Bioinformatics Core, the Flow Cytometry core at the MGH Center for Regenerative Medicine, and the MGH Center for Skeletal Research for technical assistance. M.G. was supported by the Human Frontiers Science Program Fellowship LT 000445/2015. D.T.M. was supported by F32HL154638 NIH and J.R. was supported by HHMI Faculty Scholar Award, T.D.C. was supported by NIAMS/NIH R01070139 and R21AR071554. H.L.D. was supported by the NSF GRFP. S.T, K.Z., J.L.G. were supported by AR071554 NIAMS/NIH, American Federation for Aging Research, AR072294 NIAMS/NIH and the Harvard Stem Cell Institute

## Author Contributions

Conceptualization: MG, JLG

Methodology: MG, ST, HLD, MS, KZ, MJT, MS

Investigation: MG, ST, DTM, HLD

Visualization: MG, ST, HLD

Funding acquisition: MG, JLG, TDC

Project administration: MG, JLG

Supervision: JLG

Writing – original draft: MG, ST

Writing – review & editing: MG, JLG, JR, TDC

Authors declare that they have no competing interests.

## Online Methods

### Mouse Husbandry

All mouse experiments were performed according to American Veterinary Medical Association guidelines and according to our Massachusetts General Hospital Institutional Animal Care and Use Committee (IACUC: 2013N0000062) approved protocol. The following mouse strains were used: *Axin2^tm1(cre/ERT2)Rnu^*/J (Jax Cat# 018867); H2B-GFP (*Col1a1:tetO-H2B-GFP;ROSA-rtTA* ^27^; *Scx-GFP* ^53^; *Scx-Cre^ERT2^;Porcupine^fl/fl^* (*129S-Porcn tm1.1Vdv/J,* Jax Cat# 020994); *Gt(ROSA)26Sor^tm9(CAG-tdTomato)Hze^*/J (Jax Cat# 007909); *Gt(ROSA)26Sor^tm4(ACTB-tdTomato,-EGFP)Luo^*/J (Jax Cat# 007676).

### Injuries

Excisional Achilles tendon injuries were performed using 0.3 mm biopsy punch as described^48, 54^. The incisions were sutured with 6-Ethilon nylon sutures, and tendons were harvested at specific times after injury as described^48^.

### Administration of Tamoxifen to the CreER lines

2 mg of Tamoxifen (Tam) from stock of 10mg/ml in corn oil (sigma Aldrich Cat# T5648) was given intraperitonially for 3 times on every other day at the stage described in the text (P2, P60, or 3 months of age). After the administration of Tam, we waited at least a month for the TMX to be washed out to avoid any persistent activation of CreER.

### Administration of BrdU

BrdU was injected at a concentration of 150 mg/kg (Sigma Cat#B5002) as described^20^ from the day of the injury until the collection of the tendons. For BrdU immunostaining, sections underwent antigen retrieval and immunostaining using anti-BrdU (1:100; Abcam Cat# 6326).

### H2B-GFP model

To characterize cell proliferation, we used the doxycycline (Dox) inducible histone 2B-green fluorescence protein reporter mouse H2B-GFP (*Col1a1:tetO-H2B-GFP;ROSA-rtTA)*^27^. After H2B-GFP expression is induced by Dox addition (2mg/ml; E10 to birth), Dox is removed for the chase period (starting at P0), and the GFP is diluted in proportion to cell division. Tendon cells were isolated from the hindlimbs and forelimbs at indicated stages and analyzed after negative sorting for CD45 and CD31. To determine GFP intensity, we FACS sorted specific tendon cells using GFP beads as a reference for every FACS run (molecular probes Cat# C16508) as described^20^

### RNA extraction and RT-qPCR

RNA extraction and RT-qPCR of mouse Achilles tendons were done according to^20, 48^ RT-qPCR primer sequences are as follows:

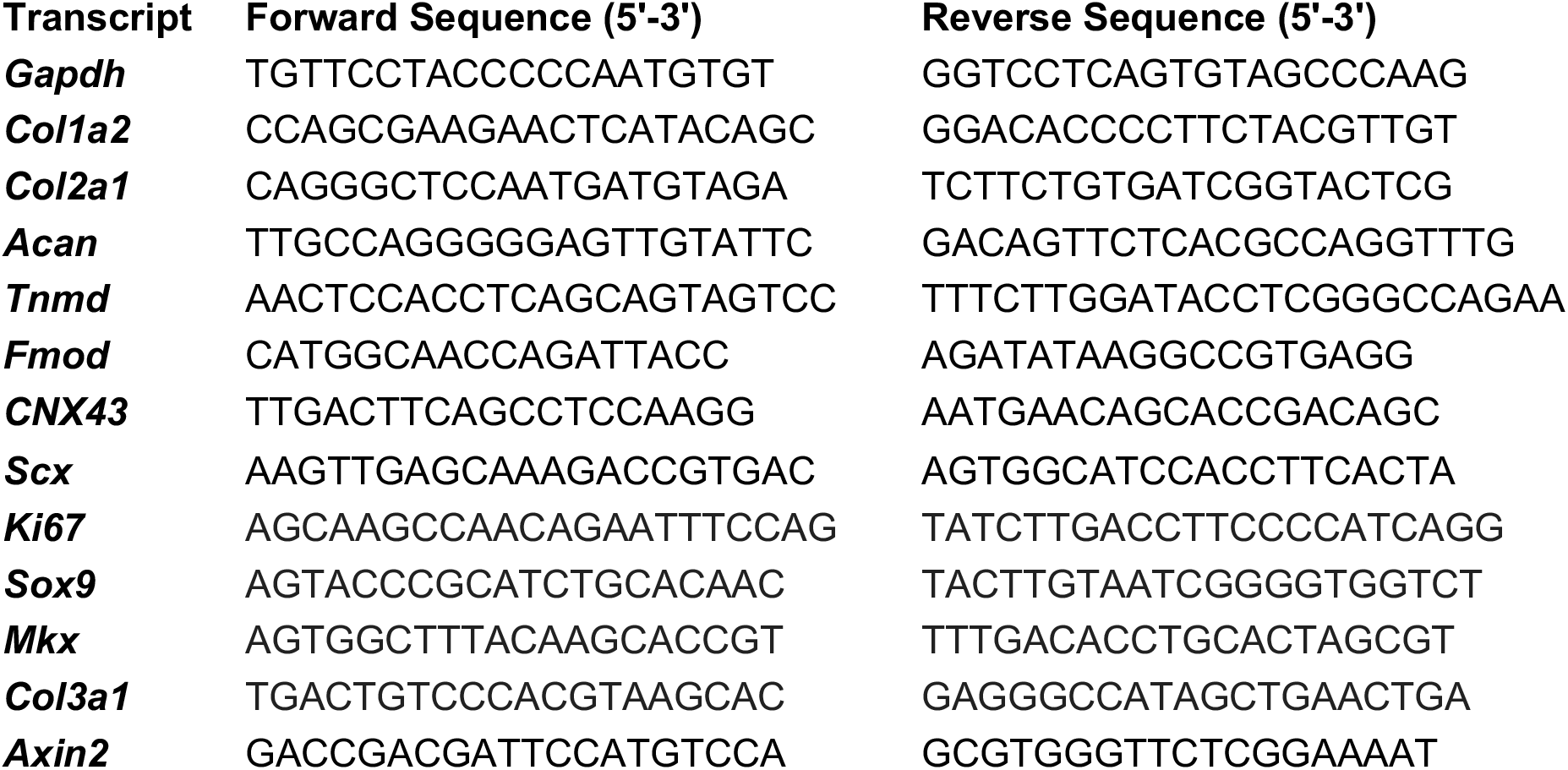

### Postnatal RNA-Seq

Extraction of intact total RNA from whole tendons was performed^20, 48^. Between 7 and 9 biological replicates (P0-8 mice, P7-7mice, P14-7 mice, P21-7 mice, P28-8 mice, P35-9 mice) per weekly time point (P0-P35) passed quality measures, yielding a total of 46 samples spanning 6 time points. RNA-seq library preparation was completed in a single batch on an Apollo 324 NGS library prep system (IntegenX) and the PrepX mRNA library protocol (Takara Bio) in the Harvard Bauer Core Facility. Single-end 75 bp reads were sequenced on an Illumina NextSeq 500 using a High-Output 75-cycle kit (Illumina) in the Harvard Bauer Core Facility. Sequenced reads were pre-processed and quantified using Salmon^55^. DESeq2 was used to determine differential expression. All downstream analyses were performed in R (R Core Team 2019).

### Postnatal ATAC-seq

*Scx*-lineage cells were isolated from *Scx-Cre;TdTom+* mice via enzymatic digestion and FACS sorted into replicates of 5,000 cells each. Transposition and library preparation were performed using a modified version of the Fast-ATAC protocol. Paired-end 42 bp reads were sequenced on an Illumina NextSeq 500 using a High-Output 75-cycle kit (Illumina) in the Harvard Bauer Core Facility. Peaks were identified from Bowtie2 aligned reads using MACS2^56^ and all downstream analyses were performed in R (R Core Team 2019) using the following packages: DiffBind^57^, ChIPseeker^58^, clusterR, ComplexHeatmap^59^, and ClusterProfiler^60^.

### Axin2^TdTom^ SMART seq

Freshly isolated tendons were collected from 4 month old *Axin2^TdTom^* vs *Axin2^-^* mouse forelimbs and hindlimbs or from injured Achilles tendons (N>10 mice; Tam at 3 months; collected at 4 months in homeostasis, or injured at 4 months and collected at 10 dpi) and dissociated as described^20, 48^. The cells were sorted using BD FACSARIA II. CD31-APC, CD45-APC Cy7 cells were excluded (BD Cat# 551262, Cat# 557659). The *Axin2^TdTom^* and the *Axin2*^-^ cells were sorted separately into 96-well plates (Eppendorf, 951020401) into 50 μl of TCL buffer (QIAGEN 1031576, supplemented with 1% β-mercaptoethanol), 200-1000 cells of *Axin2^TdTom^* / *Axin2^-^* cells were sorted into each well, and stored at −80 °C. Libraries from the cell lysates were generated using the SMART-Seq protocol^61^, with some modifications in the reverse transcription step as recently described^62^. The 96 well plates with the cell lysates were thawed on ice, spun down at 1500 RPM for 30 sec, and mixed with Agenocourt RNAclean XP SPRI beads (Beckman coulter) for RNA purification. Purified RNA was resuspended in 4 μl of Mix-1, desaturated at 72°C for 3 min and placed immediately on ice for 1 min before 7 μl of Mix-2 was added. Reverse transcription was carried out at 50°C for 90 min, followed by 5 min incubation at 85°C. 14 μl of Mix-3 were added in each well and the whole transcriptome amplification step was performed at 98 °C for 3 min, followed by 21 cycles at (98 °C for 15 s, 67 °C for 20 s, and 72 °C for 6 min), and the final extension at 72 °C for 5 min. cDNA was purified with Agencourt AMPureXP SPRI beads (Beckman coulter) as described^62^. Quality control steps were performed on samples before library construction as follows: concentration measurements using Qubit dsDNA high sensitivity assay kit; cDNA size distribution using the High-Sensitivity DNA Bioanalyzer kit. Libraries were generated using the Nextera XT library prep kit (Illumina). Combined libraries were sequenced on NextSeq 500 sequencer (Illumina), using paired-end 38 base reads, and resulted in approximately 30 million 75 bp reads per sample. Data from all samples were processed using an RNA-seq pipeline implemented in the bcbio-nextgen project (bcbio-nextgen,1.1.6a-b’2684d25’). Gene expression was quantified using Salmon (version 0.13.1) using the mm10 transcriptome (Ensembl 92, 2018-10-10). To identify differentially expressed genes in a pairwise manner, the statistical package DESeq2 (version 1.24.0) was utilized. Genes were classified as differentially expressed based on a 2-fold change in expression value and false discovery rates (FDR) below 0.05.

### Single cell RNA-seq

Achilles tendons were freshly isolated from 4 adult mice (4 months of age), 8 tendons total for homeostasis, and 3 tendons total from 3 adult mice (4 months of age) for 10 days post injury. Tendons were dissociated as described^20, 48^. Cells were sorted using BD FACSARIA II. CD31-APC, CD45-APCCy7 and TER-119/Erythroid APC-Cy7 cells were excluded (BD Cat#551262, Cat# 557659, Cat#560509). Immediately following FACS, cells were loaded to the 10X platform using Chromium next GEM single cells 3’ reagents kit V3.1. Libraries were prepared using the Single cell 3’ Reagent kit (Version 3.1 10X Genomics, Pleasanton, CA, USA). Following library preparation and quantification, libraries were sequenced by the MGH Core using NextSeq system (NextSeq 2000, HiSeq 2500) to generate 200 million reads per sample. A total of three samples were analyzed: total cells at homeostasis, total cells at 10 dpi, and tdTomato+ cells at 10 dpi from *Axin2-creERT2; flox-stop-flox-tdTomato* mice (Tam given at 3 months). Raw fastq files for each sample were first trimmed using Trimmomatic^63^ to remove low quality reads. Trimmed reads were subsequently processed using Cell Ranger and the output was analyzed using Seurat v4.0^64^. First, all three samples were individually filtered to remove cells with <100 genes, <200 UMIs, low complexity (log(genesperUMI) < 0.8), and a high mitochondrial ratio (>0.2). The samples were then normalized using the SCTransform function and clustered to identify and remove CD45+ cells that should have been removed by sorting. All three samples were then integrated together (using top 3000 variable features and top 30 principal components). PCA and UMAP dimensionality reduction was performed followed by clustering at various resolutions (a resolution of 1.0 was deemed appropriate). Major cell types present were classified into tendon, Schwann cells, skeletal muscle, pericytes, or endothelial cells based on previously characterized cell type markers.

To classify tendon cells at higher resolution, tendon cells were subset (based on Col1a1 expression) from each individual sample and then integration was performed (using top 3000 variable features and top 30 principal components). PCA and UMAP dimensionality reduction was then performed along with subsequent clustering at different resolutions (a resolution of 1.0 was ultimately determined suitable). Different tendon cell populations were then classified based on the expression of known markers for cells in different regions of the tendon during homeostasis (0 dpi) (ex. *Col22a1* for MTJ-associated tenocytes, *Pthlh* for enthesis-associated tenocytes, *Sox9* for entheseal cells etc.). Four tendon clusters that could not be readily identified based on marker expression were deemed as tendon fibroblast cells (TF1-4, see Extended data Figure 4-1A for markers). UMAP representations, dotplots and heatmaps were generated via Seurat or SCANpy^65^. Pseudotime trajectory analysis was performed on a subset of the integrated Seurat object (total homeostasis and 10 dpi samples only) using Slingshot^31^ with the root of the trajectory set to the midbody tenocyte cluster.

### Flow cytometry

Tendon cells were isolated from distal forelimb and hindlimb tendon tissue of mice from P2 - 5 months of age as indicated in the text and figure legends^20^. We enriched for tendon cells from *Axin2^TdTom^* by excluding CD31^+^ and CD45^+^ cells using FACS prior to analysis (BD Cat# 551262, Cat# 557659). DAPI staining was used to exclude dead cells. For each experiment, the gates were set using negative and positive controls for each fluorescent color that was used. For the H2B-GFP experiments, GFP beads were used (to calibrate between experiments. The following antibodies were used for flow cytometry analysis: CD90.2, CD44, SCA-1 (BD Cat# 56257, Cat# 563970, Cat# 561021), and ProCR (CD201) (Biolegend Cat# 141505).

### smFISH and Immunohistochemistry

We treated 8 µM paraffin tendon sections with TEG buffer for 5-6 hours for antigen retrieval, used proteases 3, 4 treatment step (RNA-Scope, ACD Cat#322340), and continued according to the protocol (RNAScope) for detection of the mRNA with the following probes: Hs-Axin2 (Cat# 400241); Hs-Wnt9A (Cat# 457931); Mm-Wnt9A (Cat# 405081); Mm-Axin2 (Cat# 400331); Mm,Hs-PolyA (Cat# 318631). Immunohistochemistry was performed on cryo-sections as described^20^ with anti-Sox9 (Millipore Cat# AB5535) and anti-*α*SMA (Sigma Cat# C6198).

### Cell culture

We isolated tendon cells from 4 month old mice after Tam at 3 months. After FACS, *Axin2^TdTom^* and *Axin2^-^* cells were plated together in cell culture dishes prior to analysis of surface marker expression by flow cytometry (day 10) or were plated separately for RT-qPCR analysis (day 10). DMEM (Gibco Cat# 119 65-092) with 1% P/S (Corning Cat# 30-002-cl), Fetal Bovine Serum (FBS, Gibco Cat# 97068), and 1% Hepes (Gibco Cat# 15630) was replaced every 2 days.

### Human samples

Human tendon samples were collected from patients aged 18-22 undergoing ACL reconstruction using an autograft from the semitendinosus (hamstring) tendon under IRB# 2013P001931.

### Imaging

2-Photon microscopy was used to image control and injured tendons as described^20^. To standardize the laser power “Bright Z” was adjusted. The images were taken with optical slices of 0.4 mm with 25X wet lens (XPlan N 25X WMP) on an Olympus 2P microscope FLOWVIEW FVMPE-RS. Confocal microscopy was used to image immunohistochemistry and smFISH of Achilles tendon sections using a Leica SP8 X (HC PL APO 40X/1.3 W CORR, HCX APO 63X/0.9 W U-V-I CS2). Zeiss AxioZoom V16 was used for whole mount hindlimb and Achilles tendon images and Zen software used for processing.

## Supplementary figures

**Extended data Figure 1-1.**
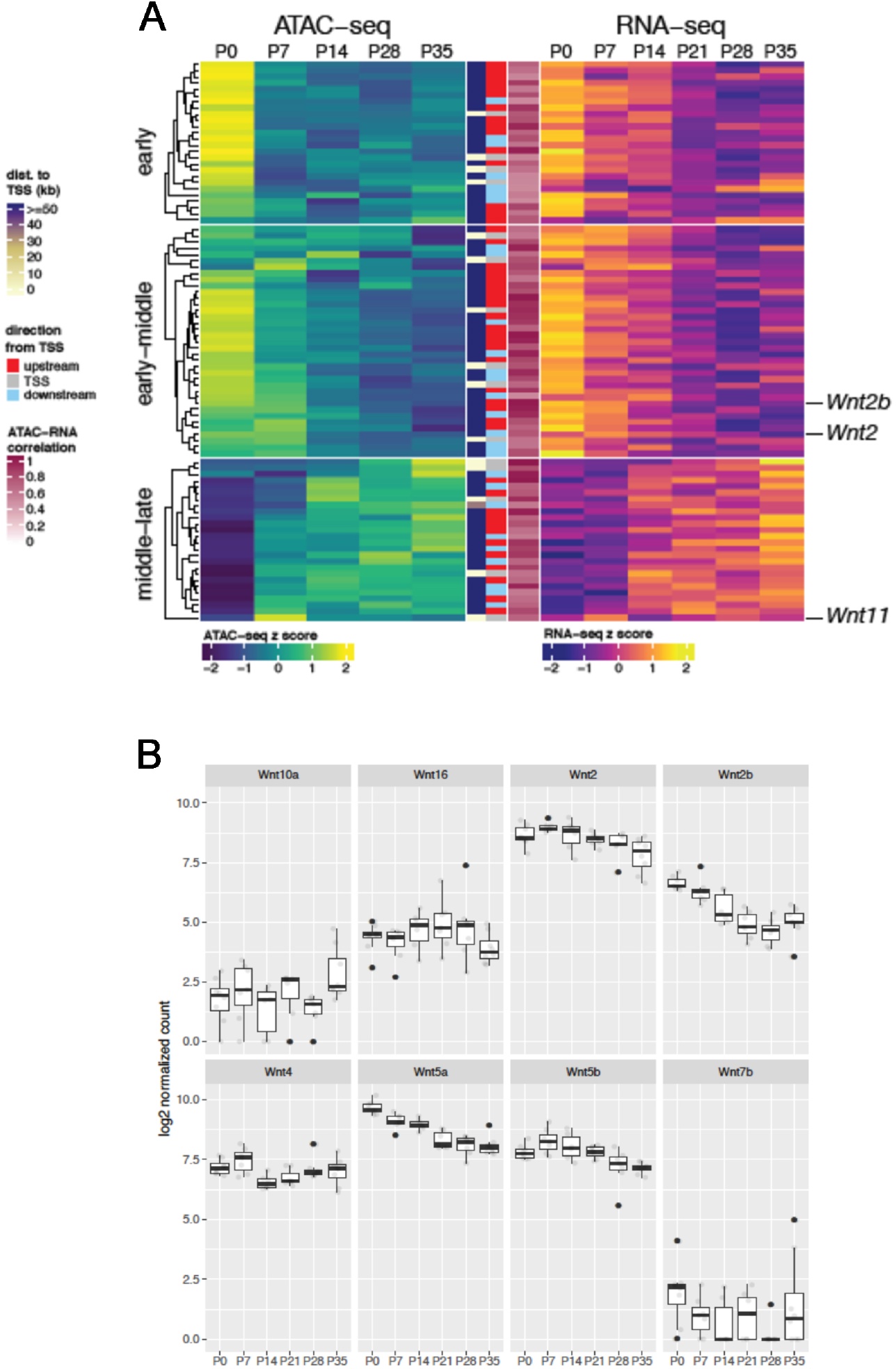
Integrated analysis of RNA-seq and ATAC-seq of distal limb tendons in postnatal stages from P0 to P35 showed an over-representation of Wnt pathway components during postnatal development (Dingwall et al., in preparation). Wnt pathway components and target genes were selected, and peaks assigned to the respective genes were extracted; non-differentially expressed genes and non-differentially accessible peaks were filtered out. After co-clustering Wnt specific peak-gene pairs, three expression and accessibility modules were identified. (Limb tendons were isolated from 7 and 9 biological replicates per weekly time point (P0-P35) for the RNAseq and 2 mice per time point for ATAC-seq. Statistics was done according to the DeSeq method as described above.) (A). Expression of Wnt ligands (*Wnt2*, *Wnt2b*, *Wnt4*, *Wnt5a*, *Wnt5b*, *Wnt7b*, *Wnt10a*, *Wnt16*) from P0-P28 in the postnatal RNA-seq. (RNA-seq analysis of Wnt genes from 7 and 9 biological replicates per weekly time point (P0-P35) presented padj< 0.05) (B).

**Extended data Figure 1-2.**
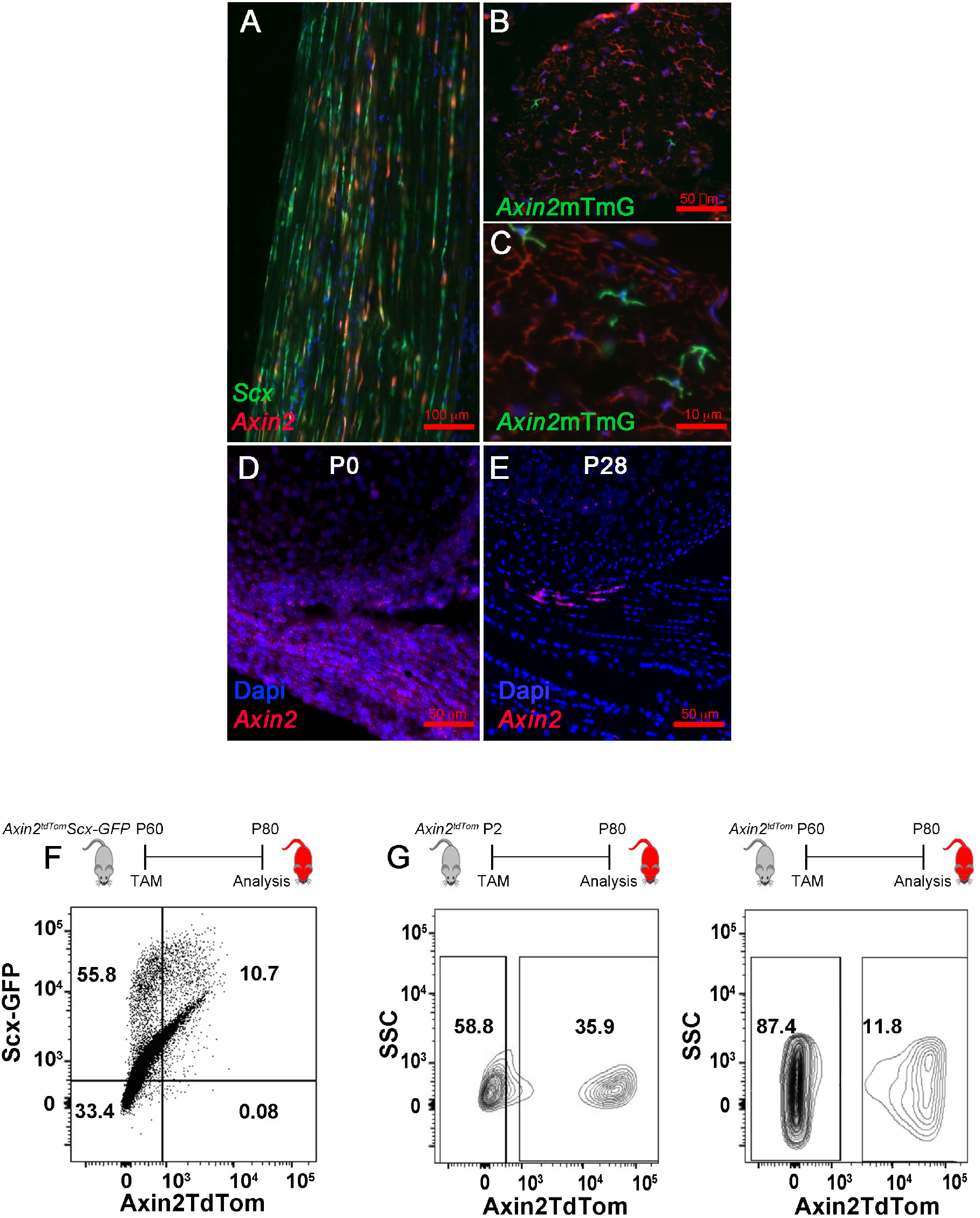
Sagittal section showing co-expressing cells in the Achilles tendon mid-body of 4-month-old *Axin2^TdTom^; Scx-GFP* mouse (Tam at 3 months) (A). Transverse sections of 4-month-old *Axin2^mTmG^* mice, showing *Axin2*-labeled cells (green) with elongated morphology in the Achilles tendon mid-body (B, C). Achilles tendon sections from n= 5 mice were analyzed (A-C). Single molecule FISH at P0 and P28 showing differing *Axin2* (red) expression in the Achilles tendon and nuclei were stained with DAPI (D, E). Representative flow cytometry plot of *Axin2^TdTom^; Scx-GFP* co-expression in adult mice from 3-5 months of age; shown is P80) (F). Representative flow cytometry plots of *Axin2^TdTom^* cell number when Tam labelling is performed at P2 vs P60 and analyzed at P80. (Achilles tendons from n>6 mice were used for each time point) (G).

**Extended data Figure 1-3 (video)**

2 Photon video of re-sliced transverse sections through an Achilles tendon of a 4 month-old mouse showing co-expression of *Axin2^TdTom^* and *Scx-GFP* and collagen matrix organization by SHG. (25X lens and 4 X digital magnification).

**Extended data Figure 3-1.**
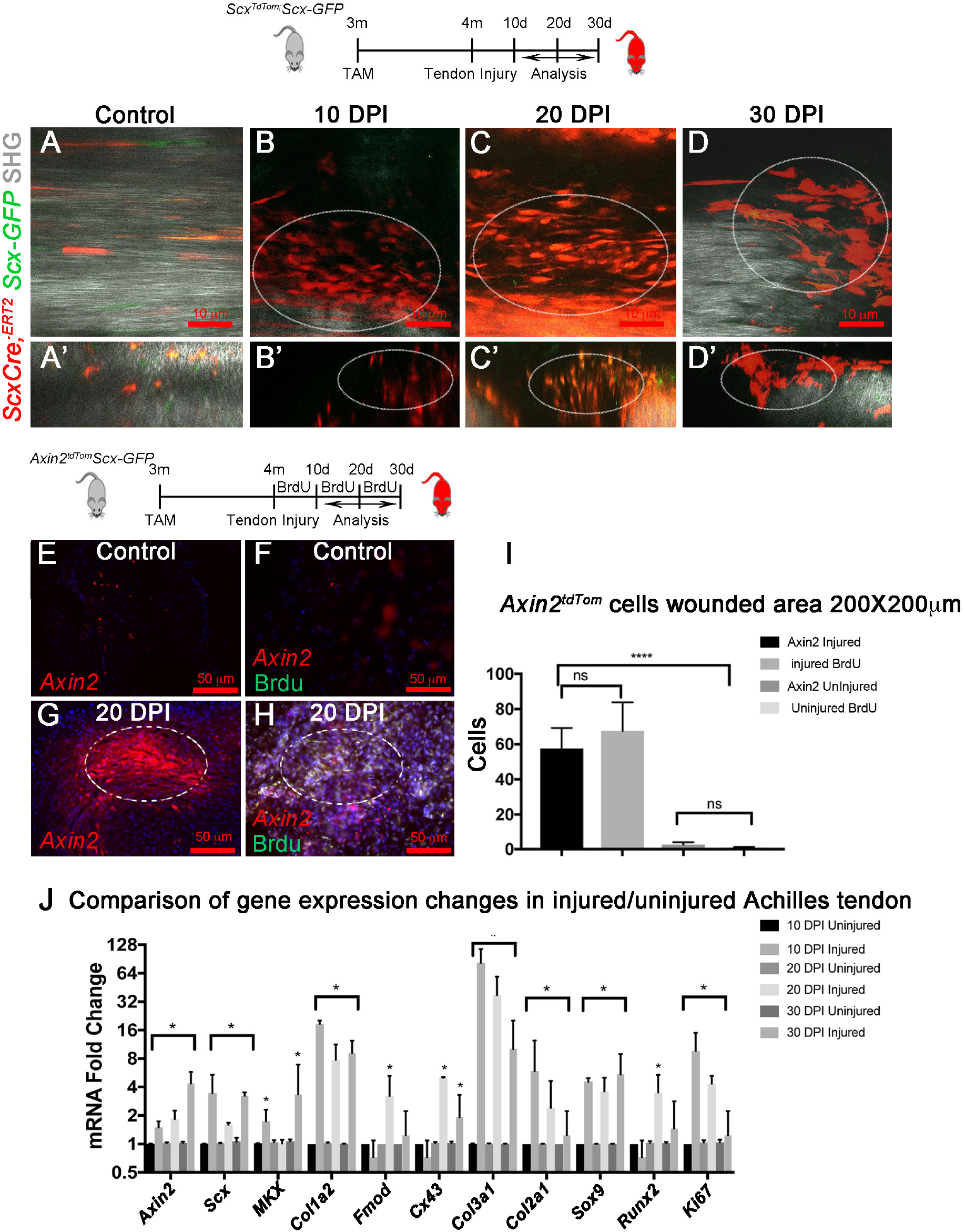
2 Photon fluorescent and SHG images of uninjured at 4 months (A) and injured Achilles tendons from *Scx^TdTom^:ScxGFP* mice at 10, 20, and 30 DPI (B-D; A-D, sagittal views; A’-D’ transverse view resliced from sagittal section of 0.4 μm). Transverse sections showing *Axin2^TdTom^* cells in uninjured controls (E) and injured tendons (G), and with BrdU labeling in uninjured at 4 months of age and injured at 20 DPI (F, H) circle indicates injured areas. Quantification of the number of *Axin2^TdTom^* cells and BrdU+ cells per defined area in injured and uninjured Achilles tendons (>10 Achilles tendon sections were analyzed per mouse with n= 44 mice and statistical analysis was calculated using one-way ANOVA; n.s.; not significant, * ****p<0.0001) (I). RT-qPCR (2^-ΔΔCT^) analysis showing increased gene expression in injured compared with uninjured contralateral control Achilles tendons at 10, 20, and 30 DPI (n=3 per time point. The multiple T-test with two-stage step-up method of Benjamini, Krieger and Yekutieli was used for statistical analysis; n.s.; not significant, *p<0.05) (J).

**Extended data Figure 3-2 (video)**

Sagittal 0.4μm sections of an Achilles tendon *Axin2^TdTom^ SCX-GFP* mouse at 10 DPI (Tam at 3 months; injury at 4 months of age) showing *Axin2^TdTom^* cells and diminished SHG signal at the injury site. (25X lens and 2 X digital magnification).

**Extended data Figure 3-3 (video)**

Sagittal 0.4μm sections of an Achilles tendon from *Axin2^TdTom^ SCX-GFP* mouse at 20 DPI (Tam at 3 months; injury at 4 months of age) showing *Axin2^TdTom^* cells co-expressing Scx-GFP and diminished SHG signal at the injury site. (25X lens and 2 X digital magnification).

**Extended data Figure 3-4 (video)**

Sagittal 0.4μm sections of an Achilles tendon from *Axin2^TdTom^ SCX-GFP* mouse at 30 DPI (Tam at 3 months; injury at 4 months of age) showing *Axin2^TdTom^* cells co-expressing *Scx-GFP* and diminished SHG signal at the injury site. (25X lens and 2 X digital magnification).

**Extended data Figure 4-1.**
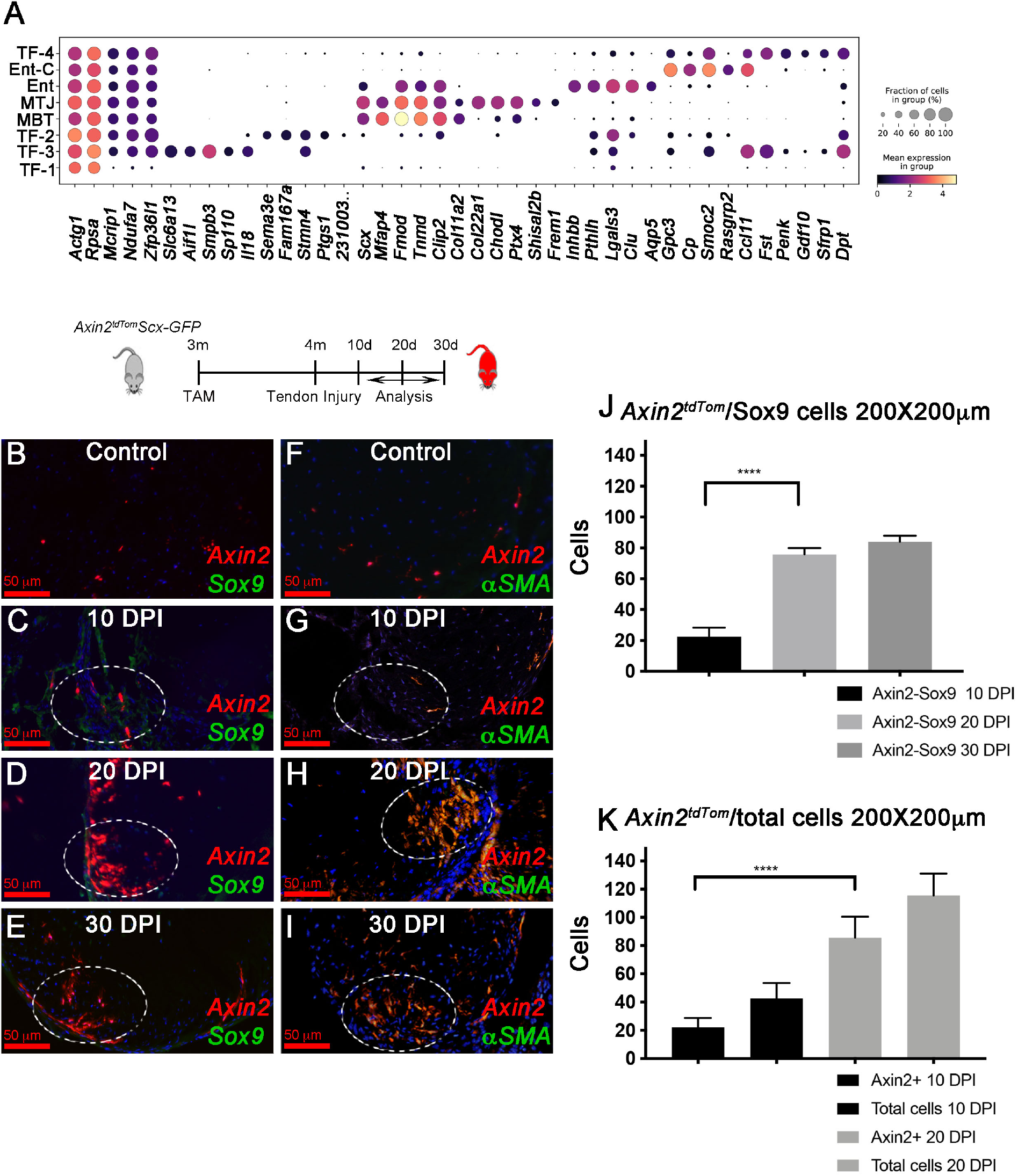
Dot plot of the most highly expressed marker genes in all of the tendon clusters during homeostasis excluding the injury-responsive cell states (A). Time course of Sox9 and αSMA co-expression with *Axin2^TdTom^* cells after injury (n> 4 mice per group; Tam given at 3 months and injury at 4 months). Transverse sections show co-expression of *Axin2^TdTom^* with Sox9 (B-E) or αSMA (F-I) at 10, 20, and 30 DPI and quantified for Sox9 (n>3 mice per group were analyzed at 10, 20, 30 DPI; in each mouse 200X200 μM square areas were examined with 3 square areas quantified per section, statistical analysis was calculated using one-way ANOVA, F=436; (****p<0.0001) (J). Quantification of total cells vs *Axin2^TdTom^* cells at 10 and 20 DPI (Achilles tendons from n>3 mice per group were quantified at 10, 20, and 30 DPI in 200 × 200 μM areas with 3 square areas quantified per section, statistical analysis was calculated using one-way ANOVA, F=1006; ****p<0.0001) (K).)

**Extended data Figure 4-2 (table)**

Excel table containing top gene markers for each tendon cluster during homeostasis excluding the injury-responsive cell states.

**Extended data Figure 5-1.**
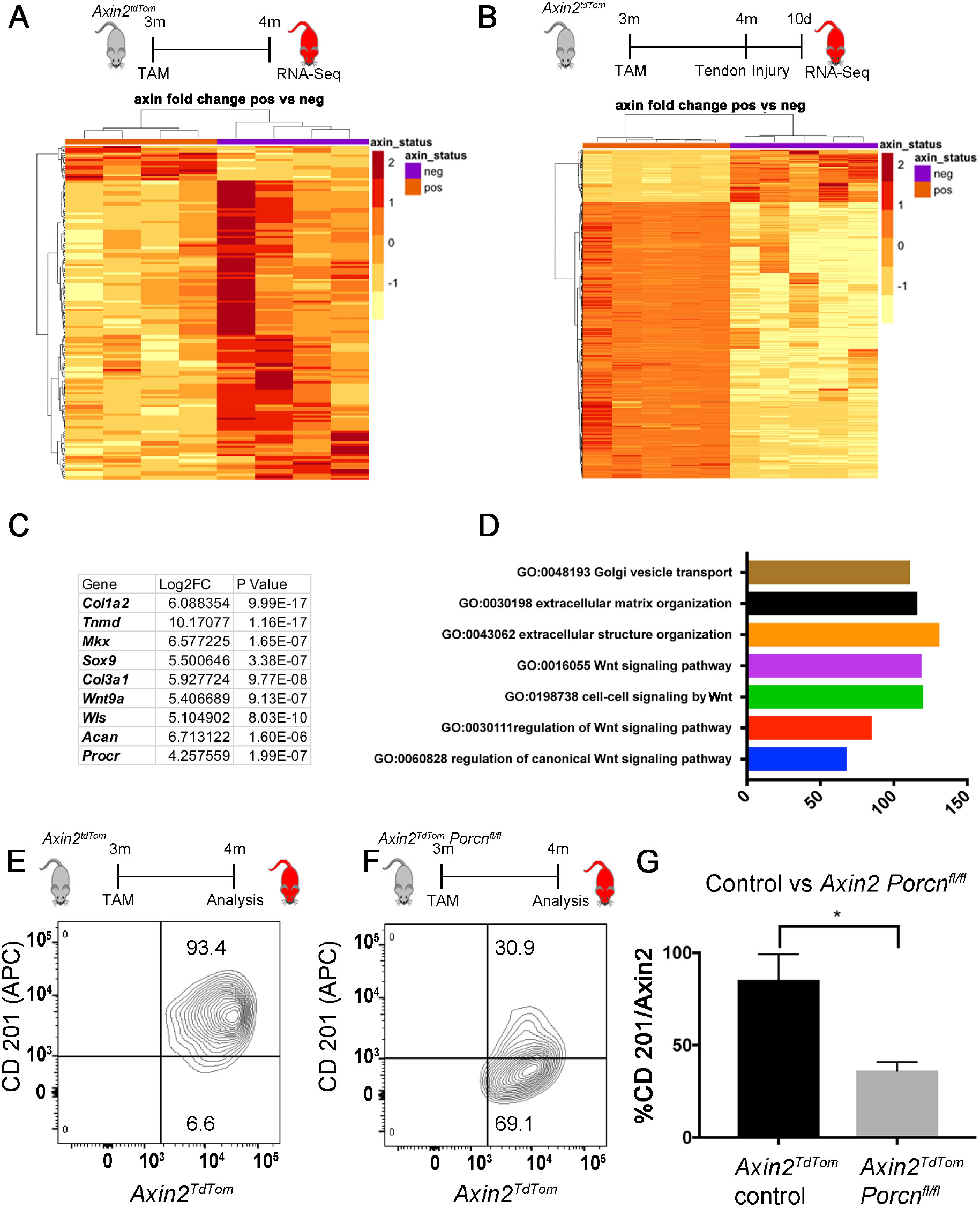
Heatmaps Axin*2^TdTom^* vs *Axin2^-^* cells in homeostasis (A) and 10 DPI (n>4 mice per group) (B). Top differentially expressed genes in the *Axin2^TdTom^* cells at 10 DPI (C). GO terms at 10 DPI shows enrichment for components of the Wnt pathway in the *Axin2^TdTom^* cells (P Value for the GO terms: GO 0048193- P Value= 9.79e-10. GO 0030198- P Value= 1.51e-4. GO 0043062- P Value 1.67e- 11. GO 0016055 P Value= 0.037. GO 0198738- P Value= 0.0346. GO 3030111- P Value= 0.00460223. GO 0060828- P Value= 0.0015406). (D). Representative FACS plots of Procr (CD201)^+^ expression in *Axin2^TdTom^* cells from wild type (E) or *Axin2:Cre^ERT2-TdTom^;Porcn^fl/fl^* tendons of 4 month old mice (F). Quantification of Procr (CD201)^+^/*Axin2^TdTom^* cells in wild type controls compared to *Axin2:Cre^ERT2-TdTom^;Porcn^fl/fl^* (Limb tendon cells were isolated from *Axin2^TdTom^* (n=3) or *Axin2-Cre^ERT2^;^TdTom^; Porcn^fl/fl^* mice (n=3) and two-way ANOVA was used for statistical analysis (F=0.508, F=25.9); *p<0.05)(G).

**Extended data Figure 6-1.**
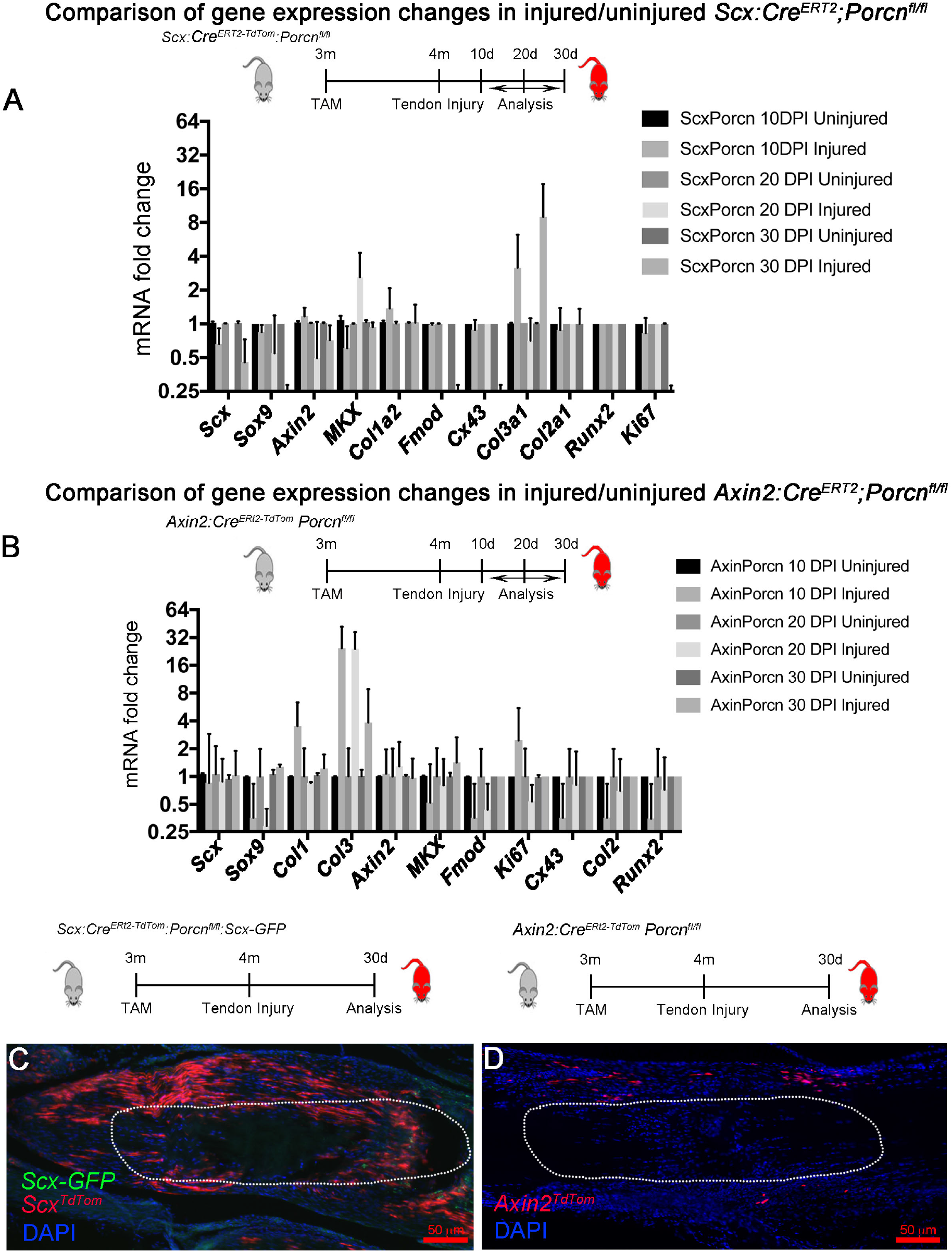
Gene expression by RT-qPCR (2^^-ΔΔCT^) is not significantly increased in injured compared with contralateral uninjured tendons of *Scx:Cre^ERT2-TdTom^;Porcn^fl/fl^* mice at 10, 20, and 30 DPI (n=3 micewere compared per time point) (A). RT-qPCR analysis shows gene expression is not significantly increased in injured compared with contralateral Achilles tendons of *Axin2:Cre^ERT2- TdTom^;Porcn^fl/fl^* at 10, 20, and 30 DPI (n=3 micewere examined per time point) (B) The multiple T-test with two-stage step-up method of Benjamini, Krieger and Yekutieli was used for statistical analysis of RT-qPCR). Sagittal sections of 5 month old *Scx:Cre^ERT2-TdTom^;Porcn^fl/fl^* (C) and *Axin2:Cre^ERT2-TdTom^;Porcn^fl/fl^* (D) at 30 DPI (n>4 mice per group) show an absence of *Axin2^TdTom^* and *Scx^TdTom^* cells at the injury site, but cells present in uninjured areas of the tendon.

**Extended data Figure 6-2.**
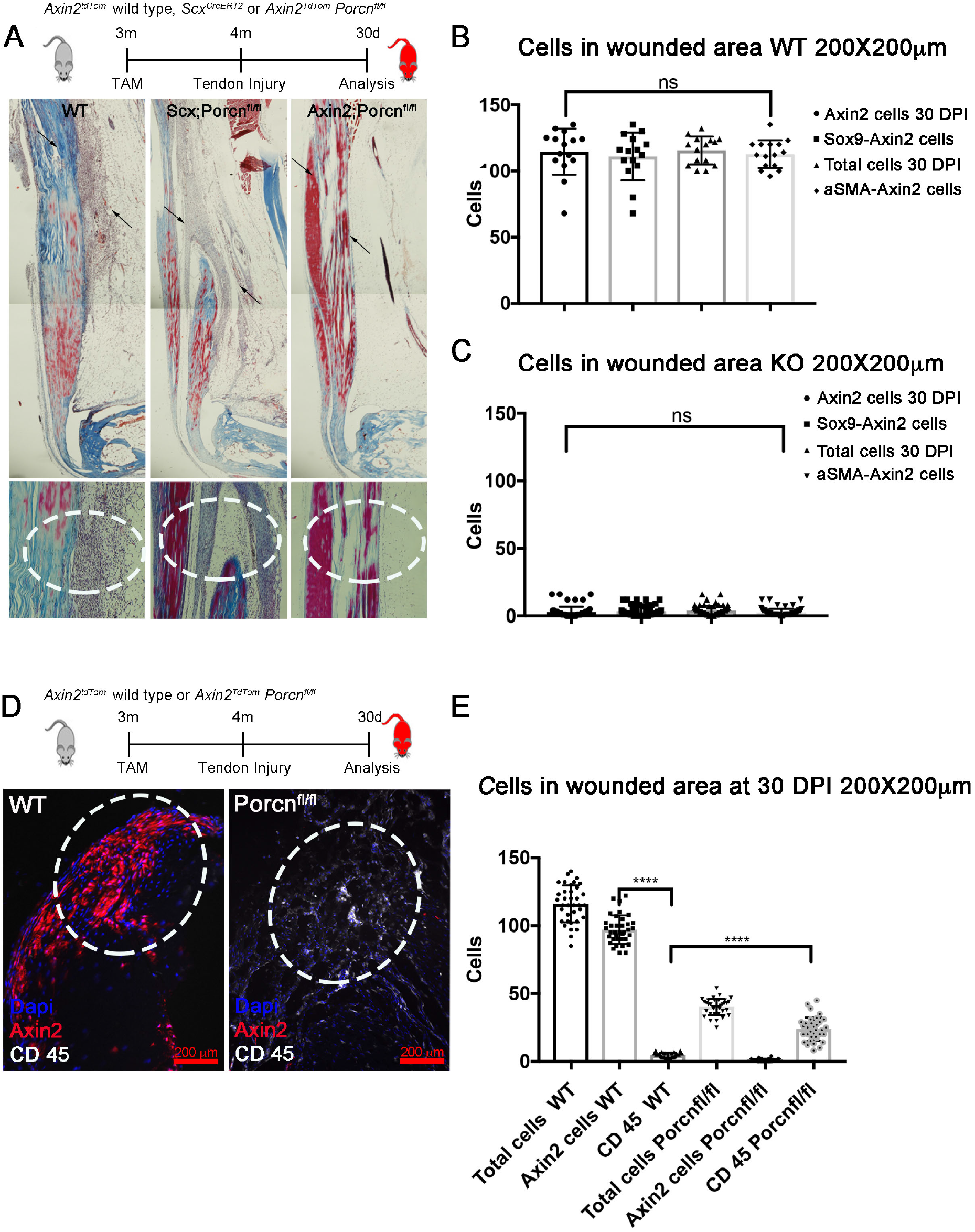
Trichome mason staining of sagittal injured Achilles tendons from wild type (WT), *Scx:Cre^ERT2^;Porcn^fl/fl^* and *Axin2:Cre^ERT2^;Porcn^fl/fl^* mice at 5 months of age. Top images show the entire Achilles tendon and injured area (arrows); bottom images are magnified views from another section of the injured area (dotted circle) (n>4 mice per group) (A). Quantification of total cells, *Axin2^TdTom^* cells, and *Axin2^TdTom^* cells co-staining with *Sox9* or *aSMA* at 30 DPI in control (B) and *Axin2:Cre^ERT2^;Porcn^fl/fl^* mutants (KO) (C) 5 month old mice (Tam at 3 months and tendon injury at 4 months; n=5 mice per group 200 ×200 μM areas were quantified with 3 squares per section and > 3 sections per mouse for each antibody; Unpaired T-test with Welch correction was used; n.s., not significant). Transverse section of injured Achilles tendon from a WT *Axin2^TdTom^* mouse (left) and *Axin2:Cre^ERT2-TdTom^;Porcn^fl/fl^* (right) mouse at 5 months of age showing increased CD45 staining in the *Axin2:Cre^ERT2-TdTom^;Porcn^fl/fl^* mutants (D). Quantification shows more CD45 cells in the *Axin2-Cre^ERT2-TdTom^; Porcn^fl/fl^* mutant injured tendons compared to WT at 30 DPI (n=4 mice per group; 200 × 200μM areas were quantified with 3 square areas per section and at least 3 sections per mouse tendon. Unpaired T test with Welch correction was used; ****p<0.0001) (E).

**Extended data Figure 6-3 (video)**

Sagittal 0.4Dm sections of an Achilles tendon from an *Axin2:Cre^ERT2-TdTom^;Porcn^fl/fl^ Scx-GFP* 5 month old mouse at 30 DPI showing absence of *Axin2^TdTom^* cells and diminished SHG signal at the injury site. (25X lens and 2 X digital magnification).

**Extended data Figure 6-4 (video)**

Sagittal 0.4Dm sections of an Achilles tendon from *Scx:Cre^ERT2-TdTom^;Porcn^fl/fl^Scx-GFP* 5 month old mouse at 30 DPI showing an absence of *Scx^TdTom^* cells and diminished SHG signal at the injury site. (25X lens and 2 X digital magnification).

